## Supplementary material for "Sibling Similarity Can Reveal Key Insights into Genetic Architecture": latex source: refs.bib

@article{rep_8,
      title={Genetic association data are broadly consistent with stabilizing selection shaping human common diseases and traits},
        author={Koch, E and Connally, N and Baya, N and Reeve, MP and Daly, M and Neale, B and Lander, ES and Bloemendal, A and Sunyaev, S},
          journal={bioRxiv},
            pages={2024--06},
              year={2024},
                publisher={Cold Spring Harbor Laboratory}
}


@article{rep_10,
      title={Trumpet plots: visualizing the relationship between allele frequency and effect size in genetic association studies},
        author={Corte, Lucia and Liou, Lathan and O’Reilly, Paul F and Garc{\'\i}a-Gonz{\'a}lez, Judit},
          journal={GigaByte},
            volume={2023},
              year={2023},
                publisher={Gigascience Press}
}


@article{non_add,
      title={Estimation of non-additive genetic variance in human complex traits from a large sample of unrelated individuals},
        author={Hivert, Valentin and Sidorenko, Julia and Rohart, Florian and Goddard, Michael E and Yang, Jian and Wray, Naomi R and Yengo, Loic and Visscher, Peter M},
          journal={The American Journal of Human Genetics},
            volume={108},
              number={5},
                pages={786--798},
                  year={2021},
                    publisher={Elsevier}
}


@article{kss,
      title={Nonparametric simultaneous tests for location and scale testing: a comparison of several methods},
        author={Marozzi, Marco},
          journal={Communications in Statistics-Simulation and Computation},
            volume={42},
              number={6},
                pages={1298--1317},
                  year={2013},
                    publisher={Taylor \& Francis}
}


@misc{sibarc,
      title = {sibArc: Software for Inference of Genetic Architecture}, 
      year={2024},  
      howpublished = {\url{http://www.sibArc.net}},
          note = {Accessed: 2024-03-30}
}


@article{how_many,
      title={Sibling variation in polygenic traits and DNA recombination mapping with UK Biobank and IVF family data},
        author={Lello, Louis and Hsu, Maximus and Widen, Erik and Raben, Timothy G},
          journal={Scientific Reports},
            volume={13},
              number={1},
                pages={376},
                  year={2023},
                    publisher={Nature Publishing Group UK London}
}


@article{lencz2021,
      title={Utility of polygenic embryo screening for disease depends on the selection strategy},
        author={Lencz, Todd and Backenroth, Daniel and Granot-Hershkovitz, Einat and Green, Adam and Gettler, Kyle and Cho, Judy H and Weissbrod, Omer and Zuk, Or and Carmi, Shai},
          journal={Elife},
            volume={10},
              pages={e64716},
                year={2021},
                  publisher={eLife Sciences Publications Limited}
}


@article{adopt1,
	  title={Family environment and the malleability of cognitive ability: A Swedish national home-reared and adopted-away cosibling
		  control study},
	    author={Kendler, Kenneth S and Turkheimer, Eric and Ohlsson, Henrik and Sundquist, Jan and Sundquist, Kristina},
	      journal={Proceedings of the National Academy of Sciences},
	        volume={112},
		  number={15},
		    pages={4612--4617},
		      year={2015},
		        publisher={National Acad Sciences}
}

@article{adopt2,
	  title={Shared environmental influences on personality: A combined twin and adoption approach},
	    author={Matteson, Lindsay K and McGue, Matt and Iacono, William G},
	      journal={Behavior Genetics},
	        volume={43},
		  pages={491--504},
		    year={2013},
		      publisher={Springer}
}


@Article{adopt3,
	  author   = {Rosa Cheesman and Avina Hunjan and Jonathan R. I. Coleman and Yasmin Ahmadzadeh and Robert Plomin and Tom A. McAdams
		  and Thalia C. Eley and Gerome Breen},
	    journal  = {Psychological Science},
	      title    = {Comparison of Adopted and Nonadopted Individuals Reveals Gene–Environment Interplay for Education in the UK
		      Biobank},
	        year     = {2020},
		  note     = {PMID: 32302253},
		    number   = {5},
		      pages    = {582-591},
		        volume   = {31}
}


@article{fullseq,
	  title={Genome-wide association studies},
	    author={Uffelmann, Emil and Huang, Qin Qin and Munung, Nchangwi Syntia and De Vries, Jantina and Okada, Yukinori and Martin,
		    Alicia R and Martin, Hilary C and Lappalainen, Tuuli and Posthuma, Danielle},
	      journal={Nature Reviews Methods Primers},
	        volume={1},
		  number={1},
		    pages={59},
		      year={2021},
		        publisher={Nature Publishing Group UK London}
}


@article{extreme,
title={Extreme-phenotype genome-wide association study (XP-GWAS): a method for identifying trait-associated variants by
sequencing pools of individuals selected from a diversity panel},
author={Yang, Jinliang and Jiang, Haiying and Yeh, Cheng-Ting and Yu, Jianming and Jeddeloh, Jeffrey A and Nettleton, Dan and
Schnable, Patrick S},
journal={The Plant Journal},
volume={84},
number={3},
pages={587--596},
year={2015},
publisher={Wiley Online Library}
}


@article{yengo2022,
title={A saturated map of common genetic variants associated with human height},
author={Yengo, Lo{\"\i}c and Vedantam, Sailaja and Marouli, Eirini and Sidorenko, Julia and Bartell, Eric and Sakaue, Saori
and Graff, Marielisa and Eliasen, Anders U and Jiang, Yunxuan and Raghavan, Sridharan and others},
journal={Nature},
volume={610},
number={7933},
pages={704--712},
year={2022},
publisher={Nature Publishing Group UK London}
}


@article{ldlgwas, 
title={Whole genome sequence analysis of blood lipid levels in> 66,000 individuals},
author={Selvaraj, Margaret Sunitha and Li, Xihao and Li, Zilin and Pampana, Akhil and Zhang, David Y and Park, Joseph and
Aslibekyan, Stella and Bis, Joshua C and Brody, Jennifer A and Cade, Brian E and others},
	      journal={Nature communications},
	        volume={13},
		  number={1},
		    pages={5995},
		      year={2022},
		        publisher={Nature Publishing Group UK London}
}


@article{schema,
	  title={Rare coding variants in ten genes confer substantial risk for schizophrenia},
	    author={Singh, Tarjinder and Poterba, Timothy and Curtis, David and Akil, Huda and Al Eissa, Mariam and Barchas, Jack D and
		    Bass, Nicholas and Bigdeli, Tim B and Breen, Gerome and Bromet, Evelyn J and others},
	      journal={Nature},
	        volume={604},
		  number={7906},
		    pages={509--516},
		      year={2022},
		        publisher={Nature Publishing Group UK London}
}


@article{newsandviews,
	author = {Iyegbe, Conrad and O’Reilly, Paul},
	year = {2022},
	month = {04},
	pages = {},
	title = {Genetic origins of schizophrenia find common ground},
	volume = {604},
	journal = {Nature},
	doi = {10.1038/d41586-022-00773-5}
}


@article{polerman,
	  title={Meta-analysis of the heritability of human traits based on fifty years of twin studies},
	    author={Polderman, Tinca JC and Benyamin, Beben and De Leeuw, Christiaan A and Sullivan, Patrick F and Van Bochoven, Arjen and
		    Visscher, Peter M and Posthuma, Danielle},
	      journal={Nature genetics},
	        volume={47},
		  number={7},
		    pages={702--709},
		      year={2015},
		        publisher={Nature Publishing Group US New York}
}


@article{wray2018,
	  title={Common disease is more complex than implied by the core gene omnigenic model},
	    author={Wray, Naomi R and Wijmenga, Cisca and Sullivan, Patrick F and Yang, Jian and Visscher, Peter M},
	      journal={Cell},
	        volume={173},
		  number={7},
		    pages={1573--1580},
		      year={2018},
		        publisher={Elsevier}
}

% Barton %
@article{barton2002,
title={Understanding quantitative genetic variation},author={Barton, Nicholas H and Keightley, Peter D},journal={Nature Reviews Genetics},volume={3},number={1},pages={11--21},year={2002},publisher={Nature Publishing Group}}
@article{barton2005,
title={Theoretical models of selection and mutation on quantitative traits},author={Johnson, Toby and Barton, Nick},journal={Philosophical Transactions of the Royal Society B: Biological Sciences},volume={360},number={1459},pages={1411--1425},year={2005},publisher={The Royal Society London}}
@article{barton2017,
title={The infinitesimal model: Definition, derivation, and implications}, author={Barton, Nicholas H and Etheridge, Alison M and V{\'e}ber, Amandine},journal={Theoretical population biology}, volume={118}, pages={50--73}, year={2017}, publisher={Elsevier}}
@article{barton2022,
title={The “New Synthesis”},author={Barton, Nicholas H},journal={Proceedings of the National Academy of Sciences},volume={119},number={30},pages={e2122147119},year={2022},publisher={National Acad Sciences}}

% Bulmer % 
@book{bulmer1980,
title={The mathematical theory of quantitative genetics.},author={Bulmer, Michael George and others},year={1980},publisher={Clarendon Press.}}
@article{bulmer1998,
title={Galton's law of ancestral heredity},author={Bulmer, Michael},journal={Heredity},volume={81},number={5},pages={579--585},year={1998},publisher={Nature Publishing Group}}


@article{jensen,
title={Sur les fonctions convexes et les in{\'e}galit{\'e}s entre les valeurs moyennes},
author={Jensen, Johan Ludwig William Valdemar},journal={Acta mathematica},volume={30},number={1},pages={175--193},year={1906},publisher={Springer}}


@article{social,
title={Personality maturation around the world: A cross-cultural examination of social-investment theory},
author={Bleidorn, Wiebke and Klimstra, Theo A and Denissen, Jaap JA and Rentfrow, Peter J and Potter, Jeff and Gosling, Samuel D},
journal={Psychological science},volume={24},number={12},pages={2530--2540},year={2013},publisher={Sage Publications Sage CA: Los Angeles, CA}}


@article{qtn,
title={Operating characteristics of the rank-based inverse normal transformation for quantitative trait analysis in genome-wide association studies},
author={McCaw, Zachary R and Lane, Jacqueline M and Saxena, Richa and Redline, Susan and Lin, Xihong},journal={Biometrics},volume={76},number={4},pages={1262--1272},year={2020},publisher={Wiley Online Library}}


@book{JKB,
title={Continuous univariate distributions, volume 2},
author={Johnson, Norman L and Kotz, Samuel and Balakrishnan, Narayanaswamy},
volume={289},
year={1995},
publisher={John wiley \& sons}
}

@article{hapgen,
title={HAPGEN2: simulation of multiple disease SNPs},author={Su, Zhan and Marchini, Jonathan and Donnelly, Peter},
journal={Bioinformatics},volume={27},number={16},pages={2304--2305},year={2011},publisher={Oxford University Press}}

@article{tgp,
title={A global reference for human genetic variation},author={1000 Genomes Project Consortium and others},journal={Nature},volume={526},number={7571},pages={68},year={2015},publisher={Nature Publishing Group}}


@article{selzam,
title={Evidence for gene-environment correlation in child feeding: Links between common genetic variation for BMI in children and parental feeding practices},author={Selzam, Saskia and McAdams, Tom A and Coleman, Jonathan RI and Carnell, Susan and O’Reilly, Paul F and Plomin, Robert and Llewellyn, Clare H},journal={PLoS genetics},volume={14},number={11},pages={e1007757},year={2018},publisher={Public Library of Science San Francisco, CA USA}}


@article{nigeria,
title={The Nigerian twin and sibling registry},author={Hur, Yoon-Mi and Kim, Jong Woo and Chung, Kee Wha and Shin, Joong Sik and Jeong, Hoe-Uk and Auta, Emmanuel},journal={Twin Research and Human Genetics},volume={16},number={1},pages={282--284},year={2013},publisher={Cambridge University Press}}


7

@article{scale,
title={Simple scaling laws control the genetic architectures of human complex traits},
author={Simons, Yuval Benjamin and Mostafavi, Hakhamanesh and Smith, Courtney Jean and Pritchard, Jonathan K and Sella, Guy},journal={bioRxiv},year={2022},publisher={Cold Spring Harbor Laboratory}}


@article{sib1,
title={Familial influences on neuroticism and education in the UK Biobank},
author={Cheesman, Rosa and Coleman, Jonathan and Rayner, Christopher and Purves, KL and Morneau-Vaillancourt, Genevieve and Glanville, Kylie and Choi, Shing W and Breen, Gerome and Eley,
Thalia C},journal={Behavior genetics},volume={50},number={2},pages={84--93},year={2020},publisher={Springer}}


@article{sib2,
title={The UK Biobank resource with deep phenotyping and genomic data},
author={Bycroft, Clare and Freeman, Colin and Petkova, Desislava and Band, Gavin and Elliott, Lloyd T and Sharp, Kevin and Motyer, Allan and Vukcevic, Damjan and Delaneau, Olivier and
O’Connell, Jared and others},journal={Nature},volume={562},number={7726},pages={203--209},year={2018},publisher={Nature Publishing Group}
}


@article{normality,
title={Power comparisons of shapiro-wilk, kolmogorov-smirnov, lilliefors and anderson-darling tests},
author={Razali, Nornadiah Mohd and Wah, Yap Bee and others},journal={Journal of statistical modeling and analytics},volume={2},number={1},pages={21--33},year={2011}}


@article{ldsc,
title={Estimation of genetic correlation via linkage disequilibrium score regression and genomic restricted maximum likelihood},
author={Ni, Guiyan and Moser, Gerhard and Ripke, Stephan and Neale, Benjamin M and Corvin, Aiden and Walters, James TR and Farh, Kai-How and Holmans, Peter A and Lee, Phil and
Bulik-Sullivan, Brendan and others},journal={The American Journal of Human Genetics},volume={102},number={6},pages={1185--1194},year={2018},publisher={Elsevier}}


@article{lush,
title={Intra-sire correlations or regressions of offspring on dam as a method of estimating heritability of characteristics},
author={Lush, Jay L},journal={Journal of animal science},volume={1940},number={1},pages={293--301},year={1940},publisher={Oxford University Press}}


@article{dwarves,
title={The genetic basis of dwarfism},
author={Francomano, Clair A},
journal={New England Journal of Medicine},volume={332},number={1},pages={58--59},year={1995},publisher={Mass Medical Soc}}

@article{giants,
title={Genetics of gigantism and acromegaly},
author={Hannah-Shmouni, Fady and Trivellin, Giampaolo and Stratakis, Constantine A},
journal={Growth hormone \& IGF research},volume={30},pages={37--41},year={2016},publisher={Elsevier}}


@article{norm2,
title={Statistical Divergences between Densities of Truncated Exponential Families with Nested Supports: Duo Bregman and Duo Jensen Divergences},
author={Nielsen, Frank},journal={Entropy},volume={24},number={3},pages={421},year={2022},publisher={MDPI}}


@book{norm1,
title={Continuous univariate distributions, volume 2},author={Johnson, Norman L and Kotz, Samuel and Balakrishnan, Narayanaswamy},volume={289},year={1995},publisher={John wiley \& sons}
}


@article{mend,
title={Genome-wide rare variant score associates with morphological subtypes of autism spectrum disorder},
author={Chan, Ada JS and Engchuan, Worrawat and Reuter, Miriam S and Wang, Zhuozhi and Thiruvahindrapuram, Bhooma and Trost, Brett and Nalpathamkalam, Thomas and Negrijn, Carol and
Lamoureux, Sylvia and Pellecchia, Giovanna and others},journal={Nature communications},volume={13},number={1},pages={1--16},year={2022},publisher={Nature Publishing Group}
}


@book{BandS,   
author={Bernardo, J. M. and Smith, A. F. M.},title={Bayesian {T}heory},address={Chichester},publisher={Wiley},year={1994}}	

@book{LandR,       
author={Little, R. J. A. and Rubin, D. B.},title={Statistical {A}nalysis with {M}issing {D}ata},address={New York},publisher={Wiley},year={1987}}


@article{turelli,
title={Genetic and statistical analyses of strong selection on polygenic traits: what, me normal?},author={Turelli, Michael and Barton, Nicholas H},
journal={Genetics},volume={138},number={3},pages={913--941},year={1994},publisher={Oxford University Press}}


@article{falconer1965,
title={The inheritance of liability to certain diseases, estimated from the incidence among relatives},author={Falconer, Douglas S},journal={Annals of human genetics},volume={29},number={1},pages={51--76},year={1965},publisher={Blackwell Publishing Ltd Oxford, UK}}
@article{falconer1967,
title={The inheritance of liability to diseases with variable age of onset, with particular reference to diabetes mellitus},author={Falconer, DS},journal={Annals of human genetics},volume={31},number={1},pages={1--20},year={1967},publisher={Wiley Online Library}}
@book{falconer1983,
title={Quantitative genetics},author={Falconer, Douglas Scott and Mackay, Trudy FC},year={1983},publisher={Longman London, UK}}


@article{fisher,
title={XV.—The correlation between relatives on the supposition of Mendelian inheritance.},
author={Fisher, Ronald A},
journal={Earth and Environmental Science Transactions of the Royal Society of Edinburgh},volume={52},number={2},pages={399--433},year={1919},publisher={Royal Society of Edinburgh Scotland Foundation}}

@book{galton,
title={Typical laws of heredity},author={Galton, Francis},year={1877},publisher={William Clowes and Sons}}


@article{wray,
title={Risk in relatives, heritability, SNP-based heritability, and genetic correlations in psychiatric disorders: a review},
author={Baselmans, Bart ML and Yengo, Lo{\"\i}c and van Rheenen, Wouter and Wray, Naomi R},journal={Biological Psychiatry},volume={89},number={1},pages={11--19},year={2021},publisher={Elsevier}}


@article{smith,
title={Heritability of liability and concordance in monozygous twins},author={Smith, Charles},journal={Annals of Human Genetics},volume={34},number={1},pages={85--91},year={1970},publisher={Wiley Online Library}}


@article{reich,
title={The use of multiple thresholds in determining the mode of transmission of semi-continuous traits},
author={Reich, Ti and James, JW and Morris, CA},journal={Annals of human genetics},volume={36},number={2},pages={163--184},year={1972},publisher={Blackwell Publishing Ltd Oxford, UK}}


% TOPIC: GENETIC ARCH %
@misc{garc1,
title={Genetic origins of schizophrenia find common ground}, author={Iyegbe, Conrad O and O’Reilly, Paul F},
year={2022}, publisher={Nature Publishing Group}}
@article{garc2,
title={An expanded view of complex traits: from polygenic to omnigenic}, author={Boyle, Evan A and Li, Yang I and Pritchard, Jonathan K},
journal={Cell},volume={169},number={7},pages={1177--1186},year={2017},publisher={Elsevier}}
@article{garc3,
title={Polygenic inheritance, GWAS, polygenic risk scores, and the search for functional variants},author={Crouch, Daniel JM and Bodmer, Walter F}, journal={Proceedings of the National Academy of Sciences},
volume={117}, number={32}, pages={18924--18933}, year={2020}, publisher={National Acad Sciences}}
@article{garc4,
title={Assessing the complex architecture of polygenic traits in diverged yeast populations},author={Cubillos, Francisco A and Billi, Eleonora and Z{\"o}rg{\"o}, Enik{\"o} and Parts, Leopold and Fargier, Patrick and Omholt, Stig and Blomberg, Anders and Warringer, Jonas and Louis,
Edward J and Liti, Gianni}, journal={Molecular ecology}, volume={20}, number={7}, pages={1401--1413}, year={2011},
publisher={Wiley Online Library}}

% RARE VARIANT SELECTION %
@article{rvs1,
title={Evolutionary evidence of the effect of rare variants on disease etiology},author={Gorlov, Ivan P and Gorlova, Olga Y and Frazier, Marsha L and Spitz, Margaret R and Amos, Christopher I},
journal={Clinical genetics},volume={79},number={3},pages={199--206},year={2011},publisher={Wiley Online Library}}

@article{rvs2,
title={Population genetics of rare variants and complex diseases},author={Maher, M Cyrus and Uricchio, Lawrence H and Torgerson, Dara G and Hernandez, Ryan D},journal={Human heredity},volume={74},number={3-4},pages={118--128},year={2012},publisher={Karger Publishers}}

@article{rvs3,
title={Evolutionary perspectives on polygenic selection, missing heritability, and GWAS},author={Uricchio, Lawrence H},journal={Human genetics},volume={139},number={1},pages={5--21},year={2020},publisher={Springer}}

@article{rvs4,
title={Unique roles of rare variants in the genetics of complex diseases in humans},author={Momozawa, Yukihide and Mizukami, Keijiro}, journal={Journal of human genetics}, volume={66}, number={1},
pages={11--23}, year={2021}, publisher={Nature Publishing Group}}

@article{gwas,
title={Chapter 11: Genome-wide association studies},author={Bush, William S and Moore, Jason H},journal={PLoS computational biology},volume={8},number={12},pages={e1002822},year={2012},publisher={Public Library of Science San Francisco, USA}}

@article{wgs,
title={Uncovering the roles of rare variants in common disease through whole-genome sequencing},author={Cirulli, Elizabeth T and Goldstein, David B},journal={Nature Reviews Genetics},volume={11},number={6},pages={415--425},year={2010},publisher={Nature Publishing Group}}


@article{sibreg,
title={The Nigerian twin and sibling registry},author={Hur, Yoon-Mi and Kim, Jong Woo and Chung, Kee Wha and Shin, Joong Sik and Jeong, Hoe-Uk and Auta, Emmanuel},
journal={Twin Research and Human Genetics},volume={16},number={1},pages={282--284},year={2013},publisher={Cambridge University Press}}


@article{twins,
title={Twin studies of brain, cognition, and behavior},author={Hewitt, John K},journal={Neuroscience and biobehavioral reviews},volume={115},pages={64},year={2020},publisher={NIH Public Access}}


@article{avi,
title={Discontinuity in the genetic and environmental causes of the intellectual disability spectrum},
author={Reichenberg, Abraham and Cederl{\"o}f, Martin and McMillan, Andrew and Trzaskowski, Maciej and Kapra, Ori and Fruchter, Eyal and Ginat, Karen and Davidson, Michael and Weiser, Mark
and Larsson, Henrik and others}, journal={Proceedings of the National Academy of sciences},volume={113}, number={4}, pages={1098--1103}, year={2016}, publisher={National Acad Sciences}}

@article{height,
title={Common variants show predicted polygenic effects on height in the tails of the distribution, except in extremely short individuals},
author={Chan, Yingleong and Holmen, Oddgeir L and Dauber, Andrew and Vatten, Lars and Havulinna, Aki S and Skorpen, Frank and Kval{\o}y, Kirsti and Silander, Kaisa and Nguyen, Thutrang T
and Willer, Cristen and others}, journal={PLoS genetics}, volume={7}, number={12}, pages={e1002439}, year={2011},
publisher={Public Library of Science San Francisco, USA}}

@article{autism,
title={Polygenic transmission disequilibrium confirms that common and rare variation act additively to create risk for autism spectrum disorders},
author={Weiner, Daniel J and Wigdor, Emilie M and Ripke, Stephan and Walters, Raymond K and Kosmicki, Jack A and Grove, Jakob and Samocha, Kaitlin E and Goldstein, Jacqueline I and Okbay,
Aysu and Bybjerg-Grauholm, Jonas and others}, journal={Nature genetics}, volume={49}, number={7}, pages={978--985},
year={2017}, publisher={Nature Publishing Group}}

@article{plomin,
title={Thinking positively: the genetics of high intelligence},journal={Intelligence}, volume={48}, pages={123--132}, year={2015}, publisher={Elsevier},
author={Shakeshaft, Nicholas G and Trzaskowski, Maciej and McMillan, Andrew and Krapohl, Eva and Simpson, Michael A and Reichenberg, Avi and Cederl{\"o}f, Martin and Larsson, Henrik and Lichtenstein,Paul and Plomin, Robert}}

@article{p_med,
title={Unveiling the genetic architecture of human disease for precision medicine}, author={Ramamoorthy, Anuradha and Yee, Sook Wah and Karnes, Jason},
journal={Clinical and Translational Science}, volume={12}, number={1}, pages={3}, year={2019}, publisher={Wiley-Blackwell}}

@misc{neale,
title = {Neale Lab Data: http://www.nealelab.is/uk-biobank/}, url = {http://www.nealelab.is/uk-biobank/}, author = {Benjamin Neale}, year={2018}}


@article{multiphen,
title={MultiPhen: joint model of multiple phenotypes can increase discovery in GWAS},journal={PloS one}, volume={7}, number={5}, pages={e34861}, year={2012},
author={O’Reilly, Paul F and Hoggart, Clive J and Pomyen, Yotsawat and Calboli, Federico CF and Elliott, Paul and Jarvelin, Marjo-Riitta and Coin, Lachlan JM},publisher={Public Library of Science San Francisco, USA}}


@article{multireview,
title={Multivariate simulation framework reveals performance of multi-trait GWAS methods}, author={Porter, Heather F and O’Reilly, Paul F},
journal={Scientific reports}, volume={7}, number={1}, pages={1--12}, year={2017}, publisher={Nature Publishing Group}}

@article{multihate,
title={PhenotypeSimulator: A comprehensive framework for simulating multi-trait, multi-locus genotype to phenotype relationships},
author={Meyer, Hannah Verena and Birney, Ewan}, journal={Bioinformatics}, volume={34}, number={17}, pages={2951--2956}, year={2018},publisher={Oxford University Press}}


@article{multievolve,
title={GeneEvolve: a fast and memory efficient forward-time simulator of realistic whole-genome sequence and SNP data},
author={Tahmasbi, Rasool and Keller, Matthew C},journal={Bioinformatics},volume={33},number={2},pages={294--296},year={2017},publisher={Oxford University Press}}


@article{rare_sitting1,
title={A phenome-wide association study of 26 mendelian genes reveals phenotypic expressivity of common and rare variants within the general population},
author={Tcheandjieu, Catherine and Aguirre, Matthew and Gustafsson, Stefan and Saha, Priyanka and Potiny, Praneetha and Haendel, Melissa and Ingelsson, Erik and Rivas, Manuel A and Priest,
James R}, journal={PLoS genetics}, volume={16}, number={11}, pages={e1008802}, year={2020}, publisher={Public Library of Science San Francisco, CA USA}}

@article{rare_sitting2,
title={Rare SLC13A1 variants associate with intervertebral disc disorder highlighting role of sulfate in disc pathology},
author={Bjornsdottir, Gyda and Stefansdottir, Lilja and Thorleifsson, Gudmar and Sulem, Patrick and Norland, Kristjan and Ferkingstad, Egil and Oddsson, Asmundur and Zink, Florian and
Lund, Sigrun H and Nawaz, Muhammad S and others}, journal={Nature communications},volume={13}, number={1}, pages={1--13}, year={2022}, publisher={Nature Publishing Group}}
@article{rare_heel,
title={Identification of 153 new loci associated with heel bone mineral density and functional involvement of GPC6 in osteoporosis},
author={Kemp, John P and Morris, John A and Medina-Gomez, Carolina and Forgetta, Vincenzo and Warrington, Nicole M and Youlten, Scott E and Zheng, Jie and Gregson, Celia L and Grundberg,
Elin and Trajanoska, Katerina and others}, journal={Nature genetics}, volume={49}, number={10}, pages={1468--1475},
year={2017}, publisher={Nature Publishing Group}}


@article{mixes,
title={Normal variance-mean mixtures and z distributions},
author={Barndorff-Nielsen, Ole and Kent, John and S{\o}rensen, Michael},
journal={International Statistical Review/Revue Internationale de Statistique},
pages={145--159},
year={1982},
publisher={JSTOR}
}


@article{order1,
title={Algorithm AS 177: Expected normal order statistics (exact and approximate)},author={Royston, JP},journal={Journal of the royal statistical society. Series C (Applied statistics)},volume={31},number={2},pages={161--165},year={1982},publisher={JSTOR}}


@article{order2,
title={On the distribution of summary statistics for missing data}, author={Ringham, BM and Kreidler, SM and Muller, KE and Glueck, DH},
journal={Communications in Statistics-Theory and Methods}, volume={48}, number={5}, pages={1149--1165}, year={2019}, publisher={Taylor \& Francis}}

@article{prs1,
title={Polygenic risk scores: from research tools to clinical instruments}, author={Lewis, Cathryn M and Vassos, Evangelos},
journal={Genome medicine}, volume={12}, number={1}, pages={1--11}, year={2020}, publisher={BioMed Central}}

@article{prs2,
title={Modeling linkage disequilibrium increases accuracy of polygenic risk scores},author={Vilhj{\'a}lmsson, Bjarni J and Yang, Jian and Finucane, Hilary K and Gusev, Alexander and Lindstr{\"o}m, Sara and Ripke, Stephan and Genovese, Giulio and Loh, Po-Ru and Bhatia,
Gaurav and Do, Ron and others}, journal={The american journal of human genetics},volume={97}, number={4}, pages={576--592}, year={2015}, publisher={Elsevier}}

@article{gwas1,
title={Genome-wide association study of more than 40,000 bipolar disorder cases provides new insights into the underlying biology},
author={Mullins, Niamh and Forstner, Andreas J and O’Connell, Kevin S and Coombes, Brandon and Coleman, Jonathan RI and Qiao, Zhen and Als, Thomas D and Bigdeli, Tim B and B{\o}rte, Sigrid
and Bryois, Julien and others}, journal={Nature genetics}, volume={53}, number={6}, pages={817--829}, year={2021},publisher={Nature Publishing Group}}

@incollection{meosis,
title={Meiosis}, author={Alberts, Bruce and Johnson, Alexander and Lewis, Julian and Raff, Martin and Roberts, Keith and Walter, Peter},
booktitle={Molecular Biology of the Cell. 4th edition}, year={2002}, publisher={Garland Science}}

@article{ukb,
title={UK biobank: an open access resource for identifying the causes of a wide range of complex diseases of middle and old age},
author={Sudlow, Cathie and Gallacher, John and Allen, Naomi and Beral, Valerie and Burton, Paul and Danesh, John and Downey, Paul and Elliott, Paul and Green, Jane and Landray, Martin and
others}, journal={PLoS medicine}, volume={12}, number={3}, pages={e1001779}, year={2015}, publisher={Public Library of Science}}
