## Supplementary material for "Sibling Similarity Can Reveal Key Insights into Genetic Architecture": latex source: sibArcRevised.tex

\RequirePackage[2020-02-02]{latexrelease} 
\documentclass[times]{zHenriquesLab-StyleBioRxiv}
%\documentclass[9pt,lineno]{elife}

% Please give the surname of the lead author for the running footer
\usepackage{relsize}
\usepackage{multirow}
\usepackage{xcolor}
\usepackage{xargs}
\usepackage{placeins}
\usepackage{amsfonts, amsmath, amsthm, amssymb} % For math fonts, symbols and environments
\usepackage[version=4]{mhchem}
\usepackage{siunitx}
\usepackage{xcolor}
\usepackage{placeins}
\usepackage[normalem]{ulem}
\usepackage[colorinlistoftodos,prependcaption,textsize=tiny]{todonotes}
\usepackage{environ}
\usepackage{etoolbox}

\DeclareCaptionType[fileext=ext]{Supplementary-Fig}
\DeclareCaptionFont{xipt}{\fontsize{11}{13}\mdseries}

\definecolor{sunyGray}{RGB}{131,134,135}                                                          
\definecolor{sunyCyan}{RGB}{0,148,254}                                                            
\newcommandx{\cyansub}[1]{\large\textcolor{sunyCyan} \noindent \textcolor{sunyCyan} {\noindent\textbf{#1}}}                             

% The toggles are initially false
\newtoggle{showFig}
\togglefalse{showFig}
\toggletrue{showFig}
%\environfinalcode{}
\NewEnviron{maybe}[1]{\iftoggle{#1}{\BODY}{}}

\newcommandx{\cnote}[2][1=]{\todo[linecolor=red,backgroundcolor=red!25,bordercolor=red,#1]{Clive: #2}}
\newcommandx{\pnote}[2][1=]{\todo[linecolor=blue,backgroundcolor=blue!25,bordercolor=blue,#1]{Paul: #2}}
\newcommandx{\tnote}[2][1=]{\todo[linecolor=green,backgroundcolor=green!25,bordercolor=green,#1]{Tade: #2}}

\newcommand{\stkout}[1]{\ifmmode\text{\sout{\ensuremath{#1}}}\else\sout{#1}\fi}
\newcommand{\Ga}{\mbox{Ga}}
\newcommand{\N}{\mathcal{N}}
%\newcommand{\L}{\mathcal{L}}
\newcommand{\E}{\mbox{E}}
\newcommand{\V}{\mbox{V}}
\newcommand{\Wi}{\mbox{Wi}}
\newcommand{\NEG}{\mbox{NEG}}
\newcommand{\Ep}{\mbox{Ep}}
\newcommand\eref[1]{{\textbf{\textit{Equation \ref{#1}}}}{}}
\newcommand\erefs[2]{\textbf{\textit{Equations \ref{#1} \& \ref{#2}}}}
\newcommand\fref[1]{\textbf{\textit{Figure \ref{#1}}}}
\newcommand{\myemph}[1]{\textbf{\textit{#1}}}
\setlength\parindent{0.50cm}
\leadauthor{Souaiaia} 
\begin{document}
\title{Sibling Similarity Can Reveal Key Insights into Genetic Architecture}
\shorttitle{Sibling Architecture}


%\mainpoint{Results of normalization}  Kolmogorov 
% Use letters for affiliations, numbers to show equal authorship (if applicable) and to indicate the corresponding author
\author[1,\Letter]{Tade Souaiaia}
\author[2]{Hei Man Wu} 
\author[2,\Letter,*]{Clive Hoggart} 
\author[2,\Letter,*]{Paul O'Reilly} 
\affil[1]{Department of Cellular Biology, Suny Downstate Health Sciences, Brooklyn, NY, USA} 
\affil[2]{Department of Genetics and Genomic Sciences, Icahn School of Medicine, Mount Sinai, 1 Gustave L Levy Pl, NY, NY, USA} 
\affil[*]{These authors contributed equally to this work} 
\onecolumn \maketitle
\vspace*{-5mm} 
\begin{corrauthor} \texttt{\small tade.souaiaia\at downstate.edu, clive.hoggart\at mssm.edu, paul.oreilly\at mssm.edu} \end{corrauthor}
\noindent\textcolor{sunyGray}{\rule{17.25cm}{1mm}}
\begin{abstract} %\normalsize
%\vspace*{1mm} 

\noindent\cyansub{Abstract}$\,$ The use of siblings to infer the factors influencing complex traits has been a cornerstone of quantitative genetics. 
Here we utilise siblings for a novel application: the 
inference of genetic architecture, specifically that relating to
individuals with extreme trait values (e.g. in the top
1\%). Inferring the genetic architecture most relevant to this group
of individuals is important because they are at greatest risk of
disease and may be more likely to harbour rare variants of large
effect due to natural selection.  We develop a theoretical framework
that derives expected distributions of sibling trait values based on an
index sibling’s trait value, estimated trait
heritability, and null assumptions that include infinitesimal genetic
effects and environmental factors that are either controlled for or
have combined Gaussian effects. This framework is then used to develop
statistical tests powered to distinguish between trait tails
characterised by common polygenic architecture from those that
include substantial enrichments of \emph{de novo} or rare variant
(Mendelian) architecture.  We apply our tests to UK Biobank data
here, although we note that they can be used to infer genetic
architecture in any cohort or health registry that includes siblings
and their trait values, since these tests do not use genetic
data. We describe how our approach has the potential to help 
disentangle the genetic and environmental causes of extreme trait
values, and to improve the design and power of future sequencing
studies to detect rare variants.
\end{abstract}
%\vspace*{-1mm} 
%%\begin{corrauthor} \texttt{\small tade.souaiaia\at downstate.edu, paul.oreilly\at mssm.edu} \end{corrauthor}
%\vspace*{-2mm} 
\noindent\textcolor{sunyGray}{\rule{17.25cm}{1mm}}
%\begin{keywords}
%siblings | genetic architecture | rare variants  
%\end{keywords}
%\clearpage \newpage 

%%%%%%%%%%%%%%%%%%%%%%%%%%%%%%%%%%%%%%%%%%%%%%%%%%%%%%%%%%%%%%%%%%%%%%%%%
%%%%%%%%%%%%%%%%%%%%%%%%%%%%%%%%%%%%%%%%%%%   INTRO  %%%%%%%%%%%%%%%%%%%%
%%%%%%%%%%%%%%%%%%%%%%%%%%%%%%%%%%%%%%%%%%%%%%%%%%%%%%%%%%%%%%%%%%%%%%%%%
\vspace*{-5mm}
\section*{Introduction} 
%%%%%%%%%%%%%%%%%%%%%%%%%%%%%%%%%%%%%%%%%%% INTRO  PARAGRAPH 1 %%%%%%%%%%

\large The fields of quantitative genetics and genetic epidemiology
have exploited the shared genetics and environment of siblings in many 
applications, notably to estimate heritability, using theory
developed a century ago \citep{fisher, lush}, and more recently
to infer a so-called `household effect' \citep{selzam}, which
contributes to genetic risk indirectly via correlation between
genetics and household environment.  Here we leverage sibling trait
data to infer genetic architecture
departing from polygenicity, specifically affecting the tails of the
trait distribution and consistent with enrichment in rare variants of
large effect.
%%%%%%%%%%%%%%%%%%%%%%%%%%%%%%%%%%%%%%%%%%%%%%%%%%%%%%%%%%%%%%%%%%%%%%%%%%%%%%%%%%%%%%%%%%%%%%%%%%%%%%%%%%%%%%%%%%%
% household polygenic
% distribution 
%%%%%%%%%%%%%%%%%%%%%%%%%%%%%%%%%%%%%%%%%%% INTRO  PARAGRAPH 2 %%%%%%%%%%%%%%%%%%%%%%%%%%%%%%%%%%%%%%%%%%%%%%%%% 

The genetic architecture of a complex trait is typically inferred from findings of multiple independent studies: 
genome-wide association studies (GWAS) identifying common variants \citep{yengo2022,ldlgwas},  whole exome or whole genome     
sequencing studies detecting rare variants \citep{schema}, and family sequencing studies designed to identify \emph{de novo} 
and rare Mendelian mutations \citep{newsandviews}. The relative contribution of each type of variant to trait heritability is a 
function of historical selection pressures on the trait in the population \citep{rep_8,rvs3}.                          
If selection has recently acted to increase the average value of a trait, then the lower tail of the 
trait distribution may become enriched for large effect rare variants over time 
because trait-reducing alleles will be subject to negative selection, and so those with 
large effects - most likely to `push' individuals to the lower tail - will be reduced to low frequencies \citep{rep_10}. 
Similarly, if the trait is subject to stabilising selection, then both tails of the trait distribution may be enriched for rare variant aetiology \citep{rvs4}.  
This can result in less accurate polygenic scores in the tails of the trait distribution \citep{height}, 
and can also produce dissimilarity between siblings beyond what is expected under polygenicity. For example, 
studies on the intellectual ability of sibling pairs have demonstrated similarity for average intellectual 
ability \citep{plomin}, regression-to-the-mean for siblings at the upper tail of the distribution \citep{plomin}, 
and complete discordance when one sibling is at the lower extreme tail of the distribution \citep{avi}. However, no theoretical framework has been developed to formally infer genetic architecture from sibling trait data. 

%Findings from such studies such as \emph{Reichenberg et al. 2016} \citep{avi} are are consistent with the presence of \emph{de novo} alleles 
%of large effect in the trait tails, alongside alternative explanations such as specific environmental exposures. 
%%%%%%%%%%%%%%%%%%%%%%%%%%%%%%%%%%%%%%%%%%%%%%%%%%%%%%%%%%%%%%%%%%%%%%%%%%%%%%%%%%%%%%%%%%%%%%%%%%%%%%%%%%%%%%%%%%%
%  Supplement
%  noindent limitations relative functional 
%%%%%%%%%%%%%%%%%%%%%%%%%%%%%%%%%%%%%%%%%%%   INTRO  PARAGRAPH 3 %%%%%%%%%%%%%%%%%%%%%%%%%%%%%%%%%%%%%%%%%%%%%%%%%

We introduce a novel theoretical framework that allows widely available sibling trait data in population          
cohorts and health registries to be leveraged to perform statistical tests that can infer complex 
genetic architecture in the tails of the trait distribution. These tests are powered to differentiate between polygenic, 
\emph{de novo} and Mendelian (i.e. rare variants of large effect) architectures (see \fref{fig:intro}); while 
these are simplifications of true complex architectures, our tests allow enrichments of the different architectures 
to be inferred and compared across traits. This framework establishes expectations about the trait distributions of siblings of index 
individuals with extreme trait values (e.g. the top 1\% of the trait) according to null assumptions of 
polygenicity, random mating and environmental factors that are either controlled for or have combined 
Gaussian effects. 
%%%%%%%%%%%%%%%%%%%%%%%%%%%%%%%%%%%%%%%%%%%%%%%%%%%%%%%%%%%%%%%%%%%%%%%%%%%%%%%%%%%%%%%%%%%%%%%%%%%%%%%%%%%%%%%%%%%
%
%
%%%%%%%%%%%%%%%%%%%%%%%%%%%%%%%%%%%%%%%%%%%   INTRO  PARAGRAPH 4 %%%%%%%%%%%%%%%%%%%%%%%%%%%%%%%%%%%%%%%%%%%%%%%%%
Critical to our framework is our derivation of the
"conditional-sibling trait distribution", which describes the trait
distribution for one individual given the quantile value of one or
more "index" siblings.  Our statistical framework, derivation of the
conditional-sibling trait distribution, and simulation study, allow us
to develop statistical tests to infer genetic architecture from data on
siblings \citep{sibreg} without the need for genetic data
(see \myemph{Model \& Methods}).  We validate the statistical power of
our tests using simulated data and apply our tests across a range of traits 
from the UK Biobank. We also release \textit{sibArc}, an open-source software tool that 
can be used to apply our tests to sibling trait data \citep{sibarc}.
Our novel framework can be extended to applications such as 
estimating heritability, characterising mating patterns, and 
inferring historical selection pressures.

%  color 
%%%%%%%%%%%%%%%%%%%%%%%%%%%%%%%%%%%%%%%%%%%   INTRO  FIGURE  %%%%%%%%%%%%%%%%%%%%%%%%%%%%%%%%%%%%%%%%%%%%%%%%%%%%%
\FloatBarrier \begin{figure}[!h] \centering \vspace*{-0.30cm}
%\begin{maybe}{showFig}\includegraphics[trim=0cm 0cm 0cm 0cm, clip,width=\linewidth]{../Figs/intro.png}\end{maybe}
\begin{maybe}{showFig}\includegraphics[trim=0cm 0cm 0cm 0cm, clip,width=16cm]{../Figs/intro.png}\end{maybe}
\caption{\textbf{Sibling Similarity Under Different Tail Architectures.}  
Left to right: When an individual's extreme trait value (top 1\%) is due to many alleles of small
effect (``Polygenic"), then their siblings' trait values are expected to show regression-to-the-mean (grey). When an individual's extreme
trait value is due to a \emph{de novo} mutation of large effect, then their siblings are expected to have trait values 
that correspond to the background distribution (green). When an individual's extreme trait value is the result of an inherited rare 
allele of large effect (``Mendelian"), then their siblings are expected to have either similarly extreme trait 
values or trait values that are drawn from the background distribution (red), depending on whether or not they inherited the same 
large effect allele.}
\label{fig:intro}\end{figure} \FloatBarrier \clearpage \newpage
%%%%%%%%%%%%%%%%%%%%%%%%%%%%%%%%%%%%%%%%%%%   END INTRO  %%%%%%%%%%%%
%  PREV: INTRO, NEXT: METHODS
%
%
%
%
%
%
% 
%
%
%
%
%
%
%
%%%%%%%%%%%%%%%%%%%%%%%%%%%%%%%%%%%%%%%%%%%%%%%%%%%%%%%%%%%%%%%%%%%%%%%%%%%%
%%%%%%%%%%%%%%%%%%%%%%%%%%%%%%%%%%%%%%%%%%%   START METHODS  %%%%%%%%%%%%%%%
%%%%%%%%%%%%%%%%%%%%%%%%%%%%%%%%%%%%%%%%%%%%%%%%%%%%%%%%%%%%%%%%%%%%%%%%%%%%
%\clearpage \newpage \section{Model and Methods} \vspace*{0.25cm} 
%\section{Model and Methods} \vspace*{0.25cm} 
\vspace*{-1cm} 
\section*{Model \& Methods} \label{sec:methods} 
\vspace*{-1.5mm} 
%%%%%%%%%%%%%%%%%%%%%%%%%%%%%%%%%%%%%%%%%%% METHODS PARAGRAPH 1 %%%%%%%%%%%%
Here we outline a framework that models the \emph{conditional-sibling trait distribution}, which describes an individual's trait 
distribution conditioned on an \emph{index} sibling's trait value.  In this section, we describe our derivation of the distribution for a 
completely polygenic trait, outline development of statistical tests that infer enrichments of rare genetic 
architectures via departures from polygenicity, and describe the simulation study used to validate and benchmark our results.  
Full derivations that complement this high-level summary of our methodology, as well as extended results from application of 
our tests to UK Biobank data, are included in the \myemph{Appendices}. 

%%%%%%%%%%%%%%%%%%%%%%%%%%%%%%%%%%%%%%%%%%%%%%%%%%%%%%%%%%%%%%%%%%%%%%%%%%%%%%%%%%%%%%%%%%%%%%%%%%%%%%%%%%%%%%%%%%
%%%%%%%%%%%%%%%%%%%%%%%%%%%%%%%%%%%%%%%%%%%   METHODS SUBSECTION-1 %%%%%%%%%%%%%%%%%%%%%%%%%%%%%%%%%%%%%%%%%%%%%%%
%%%%%%%%%%%%%%%%%%%%%%%%%%%%%%%%%%%%%%%%%%%%%%%%%%%%%%%%%%%%%%%%%%%%%%%%%%%%%%%%%%%%%%%%%%%%%%%%%%%%%%%%%%%%%%%%%%
\vspace*{-2.0mm} 
\subsection*{Conditional-Sibling Inference Framework} \label{methods:framework} ~\\
\vspace*{-3.5mm} 

\noindent Assuming completely polygenic (i.e. infinitesimal) architecture, siblings of
individuals with extreme trait values are expected to be less
extreme. This regression-to-the-mean can be understood through the two
factors \citep{falconer1983, meosis} that determine inherited genetic
variation: \textbf{(i)} average genetic contribution of
the parents (\emph{mid-parent}) to the trait value, and,
\textbf{(ii)} random genetic reassortment occurring during
meiosis. Assuming that an individual presenting an upper-tail trait
value does so due to \emph{both} high mid-parent trait value
\emph{and} genetic reassortment favouring a higher value, then a
sibling sharing common mid-parent trait value - but subject to
independent reassortment - is likely to be less extreme. How much
less extreme can be derived by first considering the conditional
distribution that relates mid-parents and their offspring. In the
simplest case, for a completely heritable continuous polygenic trait
in a large population, the offspring trait values, $s$, are normally
distributed around their mid-parent trait value, $m$, irrespective of
selection and population structure \cite{barton2005,barton2017}
\vspace*{-1.5mm}
\begin{equation}
  p(s\mid m) = \N\left( m,\sigma_f^2\right) \label{math:halfvar}
\end{equation} 
where $\sigma_f^2$ represents within family variance, which is constant
across the population. This result does not require the trait
distribution across the population to be Gaussian. If we assume that the
population distribution is Gaussian and there is random mating and no
selection, then $\sigma_f^2=$ half the population variance
\citep{bulmer1998, barton2005, wray}.  For a trait with heritability
$h^2$, trait variance can be partitioned into genetic, $\sigma_g^2$,
and environmental, $\sigma_e^2$, contributions, such that
$\sigma^2 =\sigma_g^2+\sigma_e^2$ and
$h^2=\sigma_g^2/(\sigma_g^2+\sigma_e^2)$. Assuming mean-centred
genetic and environmental trait contributions, and a trait variance of
1 (a standard normalised trait), the distribution for offspring
conditioned on the mid-parent genetic trait value, $m_g$, is:
\vspace*{-1.0mm}

\begin{equation} p(s\mid m_g) = \N\left(m_g,\sigma_g^2/2+\sigma_e^2\right)= \N\left( m_g, 1 - \frac{h^2}{2} \right)
\label{math:SgivenM} \end{equation} 

\noindent Since $p(m_g)=\N(0,h^2/2)$, from Bayes' Theorem it can be
shown that the mid-parent genetic trait value
conditional on offspring trait value, $s$, is distributed as follows:
\vspace*{-1.0mm} 

\begin{equation} p(m_g \mid s ) = \N\left(\frac{1}{2}sh^2, \frac{h^2}{2}\left(1-\frac{h^2}{2} \right) \right) 
\label{math:MgivenS} \end{equation} 

\noindent From \erefs{math:SgivenM}{math:MgivenS}, the sibling trait distribution conditional on an index sibling can be calculated as:
\vspace*{-2.5mm} 

\begin{alignat}{2} 
&p(s_2 \mid s_1 ) &&= \int p( s_2 \mid m_g ) p(m_g \mid s_1) \ dm_g \nonumber\\
&  &&=\N\left(\frac{1}{2}s_1h^2, 1-\frac{h^4}{4}\right) \label{math:S2givenS1}
\end{alignat} 

\noindent A full derivation is provided on Appendix 1. This
derivation via mid-parents can be generalised to cases where
assumptions of random mating, Gaussian population trait distribution
and no selection do not hold. When these assumptions do hold, then the
conditional sibling distribution can also be derived from the joint
sibling distribution defined by the relationship matrix
\cite{Lencz2021}. In the Appendix, we use the joint sibling distribution
to derive the distribution conditional on two sibling values. From
(\ref{math:S2givenS1}), we can infer that under complete
heritability, the siblings of an individual with a standard normal
trait value of $z$, will have a mean trait value of $\frac{z}{2}$,
with variance equal to three quarters of the population variance. In
\myemph{Appendix 1}, we further generalise the result
to binary phenotypes, increasing the
utility of this framework for further applications and theoretical
development.
%%%%%%%%%%%%%%%%%%%%%%%%%%%%%%%%%%   END METHODS SUBSECTION-1 %%%%%%%%%%%%%%%%
%%%%%%%%%%%%%%%%%%%%%%%%%%%%%%%%%%%   COMPLEX FIGURE  %%%%%%%%%%%%%%%%%%%%%%%%
\FloatBarrier \begin{figure}[!h]
\begin{maybe}{showFig} \includegraphics[trim=0cm 0cm 0cm 0cm, clip, width=\linewidth]{../Figs/side.png} \end{maybe} 
\caption{\textbf{Identifying Complex Tail Genetic Architecture.}
Conditional-sibling z-values plotted against index-sibling quantiles. Grey depicts complete polygenic architecture across 
index-sibling values. In the lower tail, an extreme scenario of \emph{de novo} architecture is shown in green, resulting in sibling
discordance. In the \emph{Upper Tail}, extreme Mendelian architecture is shown in red, whereby siblings are half 
concordant and half discordant, resulting in a bimodal conditional sibling trait distribution. Statistical tests to infer each 
type of complex tail architecture are designed to exploit these expected trait distributions.}
\label{fig:complexArk} \end{figure} \FloatBarrier %\clearpage \newpage 
%%%%%%%%%%%%%%%%%%%%%%%%%%%%%%%%%%%   COMPLEX FIGURE  %%%%%%%%%%%%%%%%%%%%%%%%%%%%%%%%%%%%%%%%%%%%%%%%%%%%%%%%%%%%


%%%%%%%%%%%%%%%%%%%%%%%%%%%%%%%%%%%%%%%%%%%%%%%%%%%%%%%%%%%%%%%%%%%%%%%%%%%%%%%%%%%%%%%%%%%%%%%%%%%%%%%%%%%%%%%%%%%%%%%%%%%%%%%%%%%%%%%%   METHODS SUBSECTION-2 %%%%%%%%%%%%%%%%%%%%%%%%%%%%%%%%%%%%%%%%%%%%%%%%%%%%%%%%%%%%%%%%%%%%%%%%%%%%%%%%%%%%%%%%%%%%%%%%%%%%%%%%%%%%%%%%%%%%%%%%%%%%%%%%%%%%%%\vspace*{-1mm} 
\subsection*{Statistical Tests for Complex Tail Architecture} \label{methods:tests} ~\\  
\vspace*{-2mm} 

\noindent In \fref{fig:complexArk}, the strategy employed to develop statistical tests for complex tail 
architecture is depicted. Our approach corresponds to testing for deviations from the expected 
conditional-sibling trait distribution under the null hypothesis of complete polygenicity in trait tails (see above for other null 
assumptions): excess discordance is indicative of an enrichment of \emph{de novo} mutations, while excess concordance indicates an enrichment of Mendelian variants, i.e. large effect variants segregating in the population. The heritability, $h^2$, 
required to define the null distribution, is estimated by maximising the log-likelihood of the conditional-sibling trait 
distribution (\eref{math:S2givenS1}) with respect to $h^2$:
\vspace*{-1mm} 

\begin{equation} \log \mathcal{L} \propto -n\log\left(1 - \frac{h^4}{4}\right) - \frac{1}{1-\frac{h^4}{4}} \sum_{i=1}^n \left( s_{2}^{(i)} - \frac{s_1^{(i)} h^2}{2}  \right)^2 \label{math:loglike} \end{equation} 

\noindent where $s_1$ and $s_2$ represent index and conditional-sibling trait values, respectively, and $n$ is number 
of sibling pairs. This allows $h^2$ to be estimated for given quantiles of the trait distribution by restricting sibling pair 
observations to those index siblings in the quantile of interest. 
To maximise power to detect non-polygenic architecture in the tails of the trait distribution, we estimate "polygenic 
heritability", $h_p^2$, from sibling pairs for which the index sibling trait value is between the $5^{th}$ and $95^{th}$ 
percentile (labelled "Distribution Body" in \fref{fig:complexArk}).

Tests for complex architecture are then performed 
in relation to index siblings whose trait values are in the tails of the distribution (e.g. the lower and upper 1\%). 
Below, $A_q$ denotes the set of sibling pairs for which the index sibling is in quantile $q$, such that 
$s_1>\Phi^{-1}(1-q)$ and $s_1<\Phi^{-1}(q)$ for the upper and lower 
tails, respectively, where $\Phi^{-1}$ is the inverse normal cumulative distribution function.

It should be noted that our estimate of $h^2$ in \eref{math:loglike} assumes no effects of shared environment. 
Polderman and colleagues \cite{polerman} found limited contribution of shared environment for most complex traits and, 
critically, our statistical tests are robust to shared environmental effects with consistent effects 
throughout the trait distribution (see \myemph{Discussion}).


%
%
%
%
%
%
%
%%%%%%%%%%%%%%%%%%%%%%%%%%%%%%%%%%%%%%%%%%%   METHODS SUBSECTION-2A %%%%%%%%%%%%%%%%%%%%%%%%%%%%%%%%%%%%%%%%%%%%%%%%%%%%
\vspace*{3mm} 
\subsubsection*{Statistical Test for De Novo Architecture} ~\\
\vspace*{-2mm} 

\noindent For inference of \emph{de novo} architecture in the tails of the trait distribution, we 
introduce a parameter, $\alpha$, to the log-likelihood defined by the 
conditional-sibling trait distribution \eref{math:loglike}: 
\begin{equation} \log \mathcal{L} \propto \frac{1}{2(1-\frac{h^4}{4})} \sum_{i \in A_q } \left( s_{2}^{(i)} - \left(\frac{1}{2} s_{1}^{(i)} h^2 + \alpha \right) \right)^2 \end{equation} 
\noindent Values of $\alpha > 0$ in the lower tail and $\alpha < 0$ in the upper tail indicate excess 
regression-to-the-mean and, thus, high sibling discordance, consistent with an enrichment of \emph{de novo} mutations 
among the index siblings. The z-statistic of the one-sided score test for $\alpha > 0$ in the lower quantile, 
$q$, relative to the null of $\alpha=0$ is (see \myemph{Appendix 2} for derivation):
\begin{equation}z=\frac{\sum_{i\in A_q}s_2^{(i)}-\frac{1}{2}s_1^{(i)}h^2} {\sqrt{n(1-\frac{h^4}{4})}}\end{equation}	
\noindent For the upper tail test of $\alpha < 0$, the above is multiplied by -1.

%%%%%%%%%%%%%%%%%%%%%%%%%%%%%%%%%%%%%%%%%%%   METHODS SUBSECTION-2B %%%%%%%%%%%%%%%%%%%%%%%%%%%%%%%%%%%%%%%%%%%%%%%%%%%%
\vspace*{3mm} 
\subsubsection*{Statistical Test for Mendelian Architecture} ~\\
\vspace*{-2mm} 

\noindent For inference of Mendelian architecture in the tails of the trait distribution, 
we compare the observed and expected tail sibling concordance, defined by the 
number of sibling pairs for which both siblings have trait values in the tail. For each index sibling in $A_q$, we calculate 
the probability that the conditional sibling is also in $A_q$, which, for the upper tail, is given by:
\begin{equation} P\left(s_2>\Phi^{-1}(q)\mid s_1\right) = 1 - \Phi \left( \frac{\Phi^{-1}(q)  - \frac{s_1 h^2}{2}}{\sqrt{1-\frac{h^4}{4}}} \right) \end{equation}
where $\Phi$ represents the normal cumulative distribution function. Denoting the mean of 
$P(s_2^{(i)}>\Phi^{-1}(q))$ across all index siblings in $A_q$ by $\pi_o$, the expected sibling concordance 
is $n\pi_o$, where $n$ is the number of index siblings in $A_q$.  Given an observed number of concordant 
siblings, $r$, the z-statistic for a one-sided score test for excess concordance is (see \myemph{Appendix 2} 
for derivation) given by:
\begin{equation} z=\frac{r-n\pi_0}{\sqrt{n\pi_0(1-\pi_0)}} \end{equation}
%%%%%%%%%%%%%%%%%%%%%%%%%%%%%%%%%%%%%%%%%%%   END METHODS SUBSECTION-2 %%%%%%%%%%%%%%%%%%%%%%%%%%%%%%%%%%%%%%%%%%%%%%%%%%%%
%
%
%
%
%
%
%
%%%%%%%%%%%%%%%%%%%%%%%%%%%%%%%%%%%%%%%%%%%%%%%%%%%%%%%%%%%%%%%%%%%%%%%%%%%%%%%%%%%%%%%%%%%%%%%%%%%%%%%%%%%%%%%%%%%%%%%
%%%%%%%%%%%%%%%%%%%%%%%%%%%%%%%%%%%%%%%%%%%   METHODS SUBSECTION-3 %%%%%%%%%%%%%%%%%%%%%%%%%%%%%%%%%%%%%%%%%%%%%%%%%%%%
%%%%%%%%%%%%%%%%%%%%%%%%%%%%%%%%%%%%%%%%%%%%%%%%%%%%%%%%%%%%%%%%%%%%%%%%%%%%%%%%%%%%%%%%%%%%%%%%%%%%%%%%%%%%%%%%%%%%%%%
\vspace*{5mm} 
\subsection*{Simulation of Conditional Sibling Data} \label{methods:simulations} ~\\
\vspace*{-2mm} 

\noindent We perform simulations using publicly available GWAS data on
multiple traits to validate our analytical model and to benchmark our
tests for complex architecture. \fref{fig:schematic} depicts the
different stages of our simulation procedure.  We start by simulating
a "parent population" (\textbf{step A}), utilsing the
  allele frequencies and effect sizes of the first 100k SNPs from a
  trait GWAS to sample genotypes and subsequently trait values
  assuming an additive model. Next, parents are randomly paired and
  their genetic trait values averaged to produce mid-parent trait
  values (\textbf{step B}), and genotypes of two offspring
  (\eref{math:halfvar}) and corresponding genetic trait values, $G$,
  are calculated (\textbf{step C}) assuming independent reassortment
  of parental alleles and unlinked SNPs.


%%%%%%%%%%%%%%%%%%%%%%%%%%%%%%%%%%%%%%%%%%%   SCHEMATIC FIGURE  %%%%%%%%%%%%%%%%%%%%%%%%%%%%%%%%%%%%%%%%%%%%%%%%%%%%%%%%%%%%
%\clearpage\newpage\vspace*{5mm} 
\FloatBarrier \begin{figure}[!h] 
\lineskip=-\fboxrule \fbox{\begin{minipage}{\dimexpr \textwidth-2\fboxsep-2\fboxrule} \centering %\vspace*{0.25cm}
%\begin{maybe}{showFig}\includegraphics[trim=0cm 0cm 0cm 0cm,clip,width=17cm]{../Figs/schematic.png} \end{maybe} 
\begin{maybe}{showFig}\includegraphics[trim=0cm 0cm 0cm 0cm,clip,width=\linewidth]{../Figs/schematic.png} \end{maybe} 
\caption{\textbf{Simulation Schematic.} 
Publicly available GWAS allele frequency and effect size data is used to simulate parent genetic trait value (\textbf{A}). Midparent 
genetic trait value (\textbf{B}) is simulated assuming random mating.  Offspring genotype and genetic trait value (\textbf{C}) is simulated 
assuming complete recombination.  Environmental variation (\textbf{D}) is added to compare with theoretical polygenic 
conditional sibling distribution. \emph{De novo} and Mendelian rare-variant effects are simulated (\textbf{E}) 
to benchmark tests for complex architecture (\textbf{F}).}
\vspace*{0.5cm} \label{fig:schematic} \end{minipage}} \end{figure} \FloatBarrier
%%%%%%%%%%%%%%%%%%%%%%%%%%%%%%%%%% SCHEMATIC FIGURE  %%%%%%%%%%%%%%%%%%%%%%%%%%%%%%%%%%%%%%%%%%%%%%%%%%%%%%%%%%%%
\vspace*{3mm} 


In \textbf{step D}, we generate offspring trait values for different
degrees of heritability by adding a Gaussian environmental effect. For
heritability, $h^2$, offspring trait values are given by $T=hG+E$,
where $G$ is the genetic effect, standardised to have
  mean 0 and variance 1, and the environmental effect $E$ is drawn 
from a normal distribution with mean 0 and variance $(1-h^2)$.  The
simulated trait has a $\mathcal{N}(0,1)$ distribution.

In \textbf{step E}, we simulate the effect of complex
tail architecture on the conditional-sibling trait distribution.  We
assume that rare variants are sufficiently penetrant to move
individuals into the tails of the distribution, independent of their
polygenic contribution. We, thus, modify sibling trait values for
individuals already in the tails (from Step D) to minimise
perturbation of the trait distribution. We simulate \emph{de novo}
tail architecture by resampling the less extreme sibling from the
background distribution, and simulate Mendelian tail architecture by
resampling the less extreme sibling from the background distribution
with probability 0.5, and from the same tail as the extreme sibling
with probability 0.5.

%%%%%%%%%%%%%%%%%%%%%%%%%%%%%%%%%% END METHODS SUBSECTION-3 %%%%%%%%%%%%%
%
%
%
%
%
%
%
%
%
%
%%%%%%%%%%%%%%%%%%%%%%%%%%%%%%%%%%%%%%%%%%%%%%%%%%%%%%%%%%%%%%%%%%%%%%%%%%%
%%%%%%%%%%%%%%%%%%%%%%%%%%%%%%%%%%%%%%%%%%%   METHODS SUBSECTION-4 %%%%%%%%
%%%%%%%%%%%%%%%%%%%%%%%%%%%%%%%%%%%%%%%%%%%%%%%%%%%%%%%%%%%%%%%%%%%%%%%%%%%
\vspace*{3mm} 
\subsection*{Application to UK Biobank Data} \label{methods:ukb} ~\\
\vspace*{-2mm} 


\noindent The UK Biobank includes data from over 21,000 siblings
\citep{how_many}.  To apply our tests to this dataset we began by
identifing continuous traits (at least 50 unique values) with at
least 5,000 sibling pairs, as defined by kinship coefficient 0.18 -
0.35 and > 0.1\% SNPs with IBS0 to distinguish from
parent-offspring \citep{sib1, sib2}. After removing outliers with
absolute trait value > 8 standard deviations from the mean, we
removed all traits with absolute skew or excess kurtosis greater
than $0.5$ to reduce the likelihood that skewed or heavy-tailed
trait distributions impact our statistical tests. The remaining
traits were standardised using rank-based inverse normal
transformation and adjusted for age, sex, recruitment centre, batch
covariates and the first 40 principle components.  After this, to
ensure a primarily additive polygenic aetiology, we
required that traits have heritability $h_{SNP}^2>0.1$ \citep{ldsc}
and that no single SNP contribute more than 0.01 to $h^2$.

We applied our method to estimate heritability from sibling pairs
(\eref{math:loglike}) across the distribution and within the trait
body ($5^{th}$ to $95^{th}$ percentiles) on the remaining traits, and
selected the eighteen traits for which both measures exceeded 30\% for
further analysis.  For each of these eighteen traits, siblings were
randomly assigned index and conditional status and both trait tails
were tested for departures from polygenicity using our \textit{De
Novo} and Mendelian tests, as well as a general Kolmogorov-Smirnov test \citep{kss}
to identify departures from the conditional sibling distribution
assuming polygenicity (\myemph{Appendix 2}).
%%%%%%%%%%%%%%%%%%%%%%%%%%%%%%%%%%%%%%%%%% END MODEL/METHODS %%%%%%%%%%%%%%%
%%%%%%%%%%%%%%%%%%%%%%%%%%%%%%%%%%%%%%%%%% END MODEL/METHODS %%%%%%%%%%%%%%%
%
%  PREV: METHODS, NEXT: RESULTS 
%
%
%%%%%%%%%%%%%%%%%%%%%%%%%%%%%%%%%%%%%%%%%%%   START RESULTS %%%%%%%%%%%%%
\section*{Results} \label{sec:results} 
Here we illustrate the conditional-sibling trait distribution, validate 
the accuracy of our analytical model using simulation, perform power analyses for 
our statistical tests for complex genetic tail architecture (see \myemph{Model \& Methods}), 
and apply our tests to trait data on thousands of siblings from the UK Biobank. 

\FloatBarrier \begin{figure}[!h]
\begin{maybe}{showFig} \includegraphics[trim=0cm 0cm 0cm 0cm, clip, width=\linewidth]{../Figs/sibdist.png}\end{maybe} 
\caption{\textbf{Conditional-Sibling Trait Distribution under Polygenic Architecture.}  A: The conditional-sibling trait 
distribution according to \eref{math:S2givenS1} for index siblings at the $1^{st}$, $25^{th}$, $50^{th}$, $75^{th}$, and 
$99^{th}$ percentile of the standardised trait distribution, when heritability is high ($h^2 = 0.95$, in orange) and moderate
($h^2 = 0.5$, in blue).  When heritability is $0.95$ conditional-sibling expectation is almost half of the index sibling z-score, 
when heritability is $0.5$ the conditional-sibling expectation is equal to $1/4$ of the index sibling z-score. 
B: The conditional distribution transformed into rank space.  An individual whose sibling is at the $99$\% percentile 
is expected to have a trait value in the $80$\% percentile when heritability is high and in the $67$\% percentile when 
heritability is moderate.}
\label{fig:zdist} \end{figure} \FloatBarrier
%\clearpage \newpage
%%%%%%%%%%%%%%%%%%%%%%%%%%%%%%%%%%%%%%%%%%%   END RESULTS SUBSECTION-1 %%%%%%%%%%%%%%%%%%%%%%%%%%%%%%%%%%%%%%%%%%%%%%%%%%%%
%
%
%
%
%
%
%
%
% underlying %


%%%%%%%%%%%%%%%%%%%%%%%%%%%%%%%%%%%%%%%%%%%%%%%%%%%%%%%%%%%%%%%%%%%%%%%%%%%%%%%%%%%%%%%%%%%%%%%%%%%%%%%%%%%%%%%%%%
%%%%%%%%%%%%%%%%%%%%%%%%%%%%%%%%%%%%%%   RESULTS SUBSECTION-1 %%%%%%%%%%%%%%%%%%%%%%%%%%%%%%%%%%%%%%%%%%%%%%%%%%%%
%%%%%%%%%%%%%%%%%%%%%%%%%%%%%%%%%%%%%%%%%%%%%%%%%%%%%%%%%%%%%%%%%%%%%%%%%%%%%%%%%%%%%%%%%%%%%%%%%%%%%%%%%%%%%%%%%%
\vspace*{-1mm} 
\subsection*{Conditional-Sibling Trait Distribution} ~\\
\vspace*{-3mm} 

\noindent In \fref{fig:zdist}{:A} the conditional-sibling trait distribution (\eref{math:S2givenS1}) is illustrated at different index 
sibling trait values (ranked percentiles). For an almost entirely heritable polygenic trait (orange), siblings of 
individuals at the $99^{th}$ percentile ($z=2.32$) have mean z-scores approximately halfway between the population 
mean and index mean (i.e. $z=1.1$). This regression-to-the-mean is greater when trait heritability is lower (blue), 
assuming (as here) independent environmental risk among siblings.

\noindent In \fref{fig:zdist}{:B} the conditional-sibling z-distribution is transformed into percentiles for interpretation in rank space. 
This distribution is skewed, especially at the tails, due to truncation at extreme quantiles (i.e. siblings cannot be 
more extreme than the top $1\%$). For a trait with $h^2 = 0.95$, siblings of individuals at the $99^{th}$ percentile (z=2.32) 
have a mean trait value at the $80^{th}$ percentile. Note that this is less extreme than the result of transforming their 
expected z-value into percentile space ($\Phi^{-1}(1.1)=86\%$), which is a consequence of Jensen's inequality \citep{jensen} given 
that the inverse cumulative distribution function of the normal is convex above zero.

We compared the theoretical conditional-sibling trait distributions to those generated from simulated 
data (see \myemph{Appendix 3}) and found that irrespective of trait used to simulate data 
(e.g. fluid intelligence, height) the two distributions did not differ significantly, 
suggesting that our analytically derived distributions are a valid model for the conditional-sibling trait distribution (\eref{math:S2givenS1}). 
\vspace*{-2mm} 
%%%%%%%%%%%%%%%%%%%%%%%%%%%%%%%%%%%%%   DIST FIGURE   %%%%%%%%%%%%%%%%%%%%%%%%%%%%%%%%%%%%%%%%%%%%%%%%%%%%%%%%%%%%
%
% sib
%
%
%%%%%%%%%%%%%%%%%%%%%%%%%%%%%%%%%%%%%%%%%%%%%%%%%%%%%%%%%%%%%%%%%%%%%%%%%%%%%%%%%%%%%%%%%%%%%%%%%%%%%%%%%%%%%%%%%%%%%%%
%%%%%%%%%%%%%%%%%%%%%%%%%%%%%%%%%%%%%%%%%%%   RESULTS SUBSECTION-2 %%%%%%%%%%%%%%%%%%%%%%%%%%%%%%%%%%%%%%%%%%%%%%%%%%%%
%%%%%%%%%%%%%%%%%%%%%%%%%%%%%%%%%%%%%%%%%%%%%%%%%%%%%%%%%%%%%%%%%%%%%%%%%%%%%%%%%%%%%%%%%%%%%%%%%%%%%%%%%%%%%%%%%%%%%%%
\subsection*{Power of Statistical Tests to Identify Complex Tail Architecture} \hfill\\ \noindent

\noindent Building on the theoretical framework introduced in the \myemph{Model \& Methods} and illustrated in the previous 
section, we develop statistical tests to identify complex architecture in the tails of the trait distribution. 
These tests leverage the fact that the similarity (or dissimilarity) in trait values among siblings provides 
information about the underlying genetic architecture (see \fref{fig:intro}). For example, high-impact \emph{de novo} 
mutations generate large dissimilarity between siblings when only one carries the unique mutant allele, while Mendelian variants 
can create excess similarity in the tails of the distribution when siblings both inherit the same mutant allele.  

%%%%%%%%%%%%%%%%%%%%%%%%%%%%%%%%%%%%%%%%%%%   POWER FIGURE  %%%%%%%%%%%%%%%%%%%%%%%%%%%%%%%%%%%%%%%%%%%%%%%%%%%%%%%%%%%%
\FloatBarrier \begin{figure}[!h] \vspace*{-0.15cm} 
\begin{maybe}{showFig} \includegraphics[trim=0cm 0cm 0cm 0cm, clip, width=\linewidth]{../Figs/simpower.png} \end{maybe} 
\caption{\textbf{Power to to detect complex tail architecture for different heritability levels, \emph{de novo} and Mendelian frequencies and sample sizes.} 
Simulation assumes highly penetrant \emph{de novo} and Mendelian frequencies of 0.05\%,0.1\%,0.2\%,0.3\%, and 0.5\%. The false-positive rate was set
at 0.05. Null simulations (red dashed line) demonstrate tests are well calibrated.}
\vspace*{0.25cm} \label{fig:simpower} \end{figure} \FloatBarrier
%%%%%%%%%%%%%%%%%%%%%%%%%%%%%%%%%%%%%%%%%%%   POWER FIGURE  %%%%%%%%%%%%%%%%%%%%%%%%%%%%%%%%%%%%%%%%%%%%%%%%%%%%%%%%%%%%
%

\noindent In \myemph{Appendix 2}, we provide detailed derivations for the statistical tests described at a high level in \myemph{Model \& Methods} 
and explain how they identify tail signatures in contrast to a polygenic background where conditional siblings 
regress-to-the-mean at a rate proportional to $h^2/2$ (\eref{math:S2givenS1}).  The performance of the two tests evaluated 
using simulated sibling data is shown in \fref{fig:simpower}.  These tests demonstrate that power to identify \emph{de novo} 
architecture is greatest when heritability is high, while power to identify Mendelian architecture is greatest when 
heritability is low. These patterns can be explained by the fact that high heritability should lead to relatively 
high similarity among siblings, and low heritability to low similarity, under polygenicity. When heritability is estimated near 50\% 
and at least 0.1\% of the population has high-impact rare aetiology, both tests are well-powered to identify each class of 
complex tail architecture. 
%%%%%%%%%%%%%%%%%%%%%%%%%%%%%%%%%%%%%%%%%%%   END RESULTS SUBSECTION-2 %%%%%%%%%%%%%%%%%%%%%%%%%%%%%%%%%%%%%%%%%%%%%%%%%%%%

%
%
%
%
%
%
%
%
%
%
% IBD
%%%%%%%%%%%%%%%%%%%%%%%%%%%%%%%%%%%%%%%%%%%%%%%%%%%%%%%%%%%%%%%%%%%%%%%%%%%%%%%%%%%%%%%%%%%%%%%%%%%%%%%%%%%%%%%%%%
%%%%%%%%%%%%%%%%%%%%%%%%%%%%%%%%%%%%%%   RESULTS SUBSECTION-3 %%%%%%%%%%%%%%%%%%%%%%%%%%%%%%%%%%%%%%%%%%%%%%%%%%%%
%%%%%%%%%%%%%%%%%%%%%%%%%%%%%%%%%%%%%%%%%%%%%%%%%%%%%%%%%%%%%%%%%%%%%%%%%%%%%%%%%%%%%%%%%%%%%%%%%%%%%%%%%%%%%%%%%%
\subsection*{Identifying Complex Tail Architecture in UK Biobank Data} \hfill\\ \vspace*{-3mm} \noindent

%%%%%%%%%%%%%%%%%%%%%%%%%%%%%%%%%%%%%%%   UKB FIGURE  %%%%%%%%%%%%%%%%%%%%%%%%%%%%%%%%%%%%%%%%%%%%%%%%%%%%%%%%%%%%
\FloatBarrier \begin{figure}[!h] \vspace*{-0.15cm}
\begin{maybe}{showFig} \includegraphics[trim=0cm 0cm 0cm 0cm, clip, width=\linewidth]{../Figs/ukb2.png} \end{maybe} 
\caption{\textbf{Analysis of Six UK Biobank Traits.} Application of
statistical tests for Mendelian and \emph{de novo} tail architecture to
sibling trait data of six UK Biobank traits. For each trait the conditional sibling mean is plotted under polygenicity (black line) 
for the heritability estimated estimated from the data. The red (high) and blue (low) 
bands represent the expected conditional sibling mean under polygenicity at different heritability values. 
Statistical tests for \emph{de novo} architecture, Mendelian architecture, and general departure from polygenicity (Kolmogorov-Smirnov Test, Dist P-val), 
were applied to conditional siblings with index siblings in the upper and lower 1\% of the distribution. 
Significant associations for the Mendelian and \emph{de novo} tests are shown in red and green respectively. 
Tail architecture that is not distinct from polygenic expectation is denoted in grey. 	
}
\label{fig:ukb} \end{figure} \FloatBarrier
%%%%%%%%%%%%%%%%%%%%%%%%%%%%%%%%%%%%%%%   UKB FIGURE  %%%%%%%%%%%%%%%%%%%%%%%%%%%%%%%%%%%%%%%%%%%%%%%%%%%%%%%%%%%%	


\noindent We applied our statistical tests for complex tail
architecture to sibling-pair data on eighteen traits from the UK
Biobank \citep{ukb}.  Here were present results from a set of six
traits with varied tail architecture: \textit{Sitting Height, Forced
Expiratory Volume, Urate, Ankle Spacing, Left Hand Grip Strength,
and Waist Circumference.}  For each trait we estimated conditional
sibling heritability via \eref{math:loglike}, and performed tests to
identify \emph{de novo} and Mendelian architecture in the lower and
upper tails of the distribution of each trait. Additionally, we also
performed a Kolmogorov-Smirnov test \citep{kss} to provide a
general test for any departures from our null model. We 
observed expected polygenic architecture in both tails for
\textit{Ankle Spacing}.  We inferred Mendelian architecture in the
lower tail for \textit{Urate}, and the upper tail for \textit{Wait
Circumference} and \textit{Left Hand Grip Strength}.  \emph{De
novo} architecture was inferred in the lower tail for
\textit{Forced Expiratory Volume} and strongly inferred in both tails 
for \textit{Sitting Height}, which is supported by evidence
from deep sequencing studies indicating that rare variants play a
substantial role in the genetic aetiology for this trait
\citep{rare_sitting1, rare_sitting2}.  In the lower tail for
\textit{Left Hand Grip Strength} we infer the presence of both
\emph{De Novo} (greater than expected mean) and Mendelian (more
concordant siblings than expected) architecture.  We note that this
could occur as a result of highly penetrant variants that are only
shared among some siblings or perhaps because at the extremes,
siblings with different handedness are unexpectedly divergent and
siblings with matching handedness are unexpectedly concordant.
Extended results for all eighteen traits analysed can be found in
\myemph{Appendix 4}.  
%%%%%%%%%%%%%%%%%%%%%%%%%%%%%%%%%%%%   END RESULTS %%%%%%%%%%%%%%%%%%%%%%%%%%%%%%%%%%%%%%%%%%%%%%%%%%%%%%%%%%%%%%%
%
%
%
%
%
%
%
%
%
%
%
%
%
%
%
%
%
%  PREV: RESULTS, NEXT: DISCUSSION
%  PREV: RESULTS, NEXT: DISCUSSION
%  PREV: RESULTS, NEXT: DISCUSSION
%  PREV: RESULTS, NEXT: DISCUSSION
%  PREV: RESULTS, NEXT: DISCUSSION
%  PREV: RESULTS, NEXT: DISCUSSION
%  PREV: RESULTS, NEXT: DISCUSSION
%  PREV: RESULTS, NEXT: DISCUSSION
%  PREV: RESULTS, NEXT: DISCUSSION
%  PREV: RESULTS, NEXT: DISCUSSION
%  PREV: RESULTS, NEXT: DISCUSSION
%  PREV: RESULTS, NEXT: DISCUSSION
%  PREV: RESULTS, NEXT: DISCUSSION
%  PREV: RESULTS, NEXT: DISCUSSION
%  PREV: RESULTS, NEXT: DISCUSSION
%  PREV: RESULTS, NEXT: DISCUSSION
%  PREV: RESULTS, NEXT: DISCUSSION
%  PREV: RESULTS, NEXT: DISCUSSION
%  PREV: RESULTS, NEXT: DISCUSSION
%  PREV: RESULTS, NEXT: DISCUSSION
%  PREV: RESULTS, NEXT: DISCUSSION
%
%
%
%
%
%
%
%
%
%
%
%
%
%
%
%
%
%%%%%%%%%%%%%%%%%%%%%%%%%%%%%%%%%%%%%%%%%%%%%%%%%%%%%%%%%%%%%%%%%%%%%%%%%%%%%%%%%%%%%%%%%%%%%%%%%%%%%%%%%%%%%%%%%
%%%%%%%%%%%%%%%%%%%%%%%%%%%%%%%%%%%%%%%%%%%   DISCUSSION - START  %%%%%%%%%%%%%%%%%%%%%%%%%%%%%%%%%%%%%%%%%%%%%%%
%%%%%%%%%%%%%%%%%%%%%%%%%%%%%%%%%%%%%%%%%%%%%%%%%%%%%%%%%%%%%%%%%%%%%%%%%%%%%%%%%%%%%%%%%%%%%%%%%%%%%%%%%%%%%%%%%
\section*{Discussion} \label{sec:discussion}


%%%%%%%%%%%%%%%%%%%%%%%%%%%%%%%%%%   DISCUSSION  PARAGRAPH 1 %%%%%%%%%%%%%%%%%%%%%%%%%%%%%%%%%%%%%%%%%%%%%%%%%%%%
In this paper, we present a novel approach to infer the genetic architecture of continuous traits, 
specifically in the tails of the distributions, from sibling trait data alone. Our approach is based 
on a theoretical framework that we develop, which derives the expected trait distributions of siblings 
conditional on the trait value of an index-sibling and the trait heritability, assuming polygenicity, random 
mating and environmental factors that are either controlled for or have combined Gaussian effects. The key 
intuition underlying the approach is that departures from the expected \emph{conditional-sibling trait 
distribution} in relation to index-siblings selected from the trait tails may be due to non-polygenic 
architecture in the tails. %\clearpage\newpage 
%%%%%%%%%%%%%%%%%%%%%%%%%%%%%%%%%%   DISCUSSION  PARAGRAPH 2 %%%%%%%%%%%%%%%%%%%%%%%%%%%%%%%%%%%%%%%%%%%%%%%%%%%%

%\noindent 
We demonstrate the validity of our conditional-sibling analytical derivations through simulations and show that 
our tests for identifying \emph{de novo} and Mendelian architecture in the tails of trait distributions are well-powered when 
large effect alleles are present in the population on the order of 1 out of 1000 individuals.  Applying our test to 
a subset of well-powered traits in the UK Biobank, we find evidence of complex genetic architecture in at least one tail 
($\alpha < 0.05$) in sixteen of eighteen traits and find \emph{de novo} architecture occurring more frequently than Mendelian 
architecture (19 vs 6 of 36 total tails). 

%%%%%%%%%%%%%%%%%%%%%%%%%%%%%%%%%%   DISCUSSION  PARAGRAPH 3 %%%%%%%%%%%%%%%%%%%%%%%%%%%%%%%%%%%%%%%%%%%%%%%%%%%%

There are several areas in which our work could have short-term utility. Firstly, those individuals inferred as having rare 
variants of large effect could be followed up in multiple ways to gain individual-level insights. For example, they could undergo clinical 
genetic testing to identify potential pathogenic variants with effects beyond the examined trait, either in the form of diseases or 
disorders that the individual has already been diagnosed with or else that they have yet to present with but may be at high 
future risk for. Furthermore, investigation of their environmental risk profile may indicate an alternative - environmental - 
explanation for their extreme trait value (see below), rather than the rare genetic architecture inferred by our tests.
%%%%%%%%%%%%%%%%%%%%%%%%%%%%%%%%%%   DISCUSSION  PARAGRAPH 4 %%%%%%%%%%%%%%%%%%%%%%%%%%%%%%%%%%%%%%%%%%%%%%%%%%%%

Our framework could also help to refine the design of sequencing studies for identifying rare variants of large effect. 
Such studies either sequence entire cohorts at relatively high cost \citep{fullseq} or else perform more targeted sequencing of individuals in 
the trait tails with the goal of optimising power per cost \citep{extreme}. However, even the latter approach is usually performed blind to evidence 
of enrichment of rare variant aetiology in the tails. Since our approach enables identification of individuals that may be most likely to 
harbour rare variants, then these individuals could be prioritised for (deep) sequencing. Moreover, our ability to distinguish 
between \emph{de novo} and Mendelian architecture could influence the broad study design, with the former suggesting that a family 
trio design may be more effective than population sequencing, which may be favoured if Mendelian architecture is inferred. Furthermore, 
our approach could be applied as a screening step to prioritise those traits, and corresponding tails, 
most likely to harbour rare variant architecture. Finally, if sequence data have already been collected, either cohort-wide or 
using a more targeted design, then our approach could be utilised to increase the power of statistical methods for detecting rare 
variants by upweighting individuals most likely to harbour rare variants. 
%%%%%%%%%%%%%%%%%%%%%%%%%%%%%%%%%%   DISCUSSION  PARAGRAPH 5 %%%%%%%%%%%%%%%

This study has several limitations. First and foremost, departures
from the expected conditional-sibling trait distributions could be due
to environmental risk factors, such as medication-use or work related
exposures, rather than rare genetic architecture. Thus,
while we believe that tail-specific deviations from polygenic
expectation are interesting whether they arise primarily from
genetic or environmental factors, we caution against
over-interpretation of our results. Rejection of the null
hypothesis from our tests should be considered only as indicating
effects \emph{consistent with} non-polygenic genetic architecture,
alongside alternative explanations such as tail-specific unshared
(\emph{de novo}) or shared ("Mendelian") environmental risks.  We
suggest that further investigation of individuals' clinical,
environmental and genetic profiles are required to achieve greater
certainty about the causes of their extreme trait
values. Nevertheless, given knowledge that rare variants of
large effect contribute to complex trait architecture, we expect
that traits for which we infer non-polygenic architecture, will, on
average, be more enriched for rare architecture in the tail(s) than
other traits. Secondly, our modelling assumes that environmental risk
factors of siblings are independent of each other.  If in fact shared
environmental risk factors contribute significantly to trait
similarity among siblings, then our heritability estimates will be
upwardly biased. However, this would only impact our tests if the
degree of shared environmental risk differed in the tails relative to
the rest of the trait distribution. Moreover, a large meta-analysis of
heritability estimates from twin studies \citep{polerman} concluded
that the contribution of shared environment among siblings (even
twins) is insubstantial, and so we might expect this to have limited
impact on results from our tests. Thirdly, our modelling assumes
random mating and so the results from our tests in relation to traits
that may be the subject of assortative mating should be considered
with caution. Likewise, our modelling assumes additivity of genetic
effects and so, while additivity is well-supported by much statistical
genetics research \citep{wray2018, non_add}, results from our tests
should be reconsidered for any traits with evidence for significant
non-additive genetic effects.
%%%%%%%%%%%%%%%%%%%%%%%%%%%%%%%%%%   DISCUSSION  PARAGRAPH 6 %%%%%%%%%%%%%%%%%%%%%%%%%%%%%%%%%%%%%%%%%%%%%%%%%%%%

Our approach not only provides a novel way of inferring genetic architecture (without genetic data) but can do so specifically 
in the tails of trait distributions, which are most likely to harbour complex genetic architecture, due to selection, and are a 
key focus in biomedical research given their enrichment for disease. This work could also have broader implications in 
quantitative genetics since we derive fundamental results about the relationship among family members' complex trait values. 
The conditional-sibling trait distribution provides a simple way of understanding the expected trait values of individuals 
according to their sibling's trait value, which could be used to answer questions of societal importance and inform future research. 
For example, it can be used to answer questions such as: as a consequence of genetics alone, how much overlap should 
there be in the traits of offspring of mid-parents at the $5^{th}$ and $95^{th}$ percentile and how does that contrast with what we observe 
in highly structured societies? Moreover, further development of the theory described here could lead to a range of other 
applications, for example, estimating levels of assortative mating, inferring historical selection pressures, and quantifying heritability 
in specific strata of the population. 


%%%%%%%%%%%%%%%%%%%%%%%  ACKNOWLEDGEMENTS AND CITATIONS %%%%%%%%%%%%%%%%%%%%%%%%%%%%%%%%%%%%%%%%%%%%
\section*{Acknowledgments}
We thank the participants in the UK Biobank and the scientists involved in the construction of this resource for making the sibling 
data used in this manuscript available. The work in this manuscript has been conducted using the UK Biobank Resource under application 
18177 (Dr O’Reilly). We would also like to thank Dr. Avi Reichenberg for early discussions and Dr. Peter Visscher for highlighting key references related to the topic, and Dr. Shai Carmi for providing feedback 
on a draft version of the article. 

%We would also like to thank Dr. Peter Visscher for highlighting some important references. 
%\nocite{*} % This command displays all refs in the bib file. PLEASE DELETE IT BEFORE YOU SUBMIT YOUR MANUSCRIPT!
\section*{References}
\bibliography{refs}
%%%%%%%%%%%%%%%%%%%%%%%  END %%%%%%%%%%%%%%%%%%%%%%%%%%%%%%%%%%%%%%%%%%%%


\clearpage \newpage
%%%%%%%%%%%%%%%%%%%%%%%%%%%%%%%%%%%%%%%%%% END DISCUSSION %%%%%%%%%%%%%%%%%%%%%%%%%%%%%%%%%%%%%%%%%%%%%%%%%%%%%%%
%
%
%
%
%
%
%
%
%
%
%
%
%  PREV: DISCUSSION, NEXT: APPENDIX-1
%  PREV: DISCUSSION, NEXT: APPENDIX-1
%  PREV: DISCUSSION, NEXT: APPENDIX-1
%  PREV: DISCUSSION, NEXT: APPENDIX-1
%  PREV: DISCUSSION, NEXT: APPENDIX-1
%  PREV: DISCUSSION, NEXT: APPENDIX-1
%  PREV: DISCUSSION, NEXT: APPENDIX-1
%  PREV: DISCUSSION, NEXT: APPENDIX-1
%  PREV: DISCUSSION, NEXT: APPENDIX-1
%  PREV: DISCUSSION, NEXT: APPENDIX-1
%  PREV: DISCUSSION, NEXT: APPENDIX-1
%  PREV: DISCUSSION, NEXT: APPENDIX-1
%  PREV: DISCUSSION, NEXT: APPENDIX-1
%  PREV: DISCUSSION, NEXT: APPENDIX-1
%  PREV: DISCUSSION, NEXT: APPENDIX-1
%
%
%
%
%
%
%
%
%
%
%  unshared
%%%%%%%%%%%%%%%%%%%%%%%%%%%%%%%%%%%%%%%%%%%%%%%%%%%%%%%%%%%%%%%%%%%%%%%%%%%%%%%%%%%%%%%%%%%%%%%%%%%%%%%%%%%%%%%%%
%%%%%%%%%%%%%%%%%%%%%%%%%%%%%%%%%%%%%%%%%% APPENDIX1 %%%%%%%%%%%%%%%%%%%%%%%%%%%%%%%%%%%%%%%%%%%%%%%%%%%%%%%%%%%%
%%%%%%%%%%%%%%%%%%%%%%%%%%%%%%%%%%%%%%%%%%%%%%%%%%%%%%%%%%%%%%%%%%%%%%%%%%%%%%%%%%%%%%%%%%%%%%%%%%%%%%%%%%%%%%%%%


\section*{Appendix 1: Derivation of Conditional Sibling Distributions} 
\label{app:part1}
Here we derive the distribution that describes the probability density
of the "conditional" sibling ($S_2$) given the genetic liability,
trait value and case-status of one or more index siblings. In each
case we assume a population of unrelated parents and rely on the
results from the \emph{infinitesimal polygenic model} that show that
within family variance is normally distributed around midparent
genetic liability (average of parents) with half the ancestral trait
variance even when selection, drift, population structure or dominance
effects alter the between family trait distribution \citep{barton2002,
  barton2017, barton2022}.
%%%%%%%%%%%%%%%%%%%%%%%%%%%%%%%%%%%%%%%%%%%%%%%%%%%%%%%%%%%%%%%%%%%%%%%%%%%%%
%%%%%%%%%%%%%%%%%%%%%%%  APPENDIX 1 - CASE 1  %%%%%%%%%%%%%%%%%%%%%%%%%%%%%%%
%%%%%%%%%%%%%%%%%%%%%%%%%%%%%%%%%%%%%%%%%%%%%%%%%%%%%%%%%%%%%%%%%%%%%%%%%%%%%
\subsection*{\normalsize Case 1) Index liability known, continuous trait ($h^2=1$): $P(S_2 \mid S_1 = {s_1})\hspace{2em}$} ~\\ 
\noindent We begin with the simplest case, a polygenic normally
distributed trait which is fully heritable ($h^2=1$) where the genetic
liability of an index sibling in a population is known.  Throughout we
denote the midparent, index sibling and conditional sibling by $M,S_1$
and $S_2$. We begin by calculating the midparent distribution
conditional on index liability using Bayes theorem:
\begin{alignat}{3}
  p(M=m \mid S=s ) &\propto p( S=s \mid M=m ) p(M=m) \nonumber \\
                   &=\N\left(S\mid m, \frac{\sigma^2}{2}\right)\N(M\mid 0, \sigma^2/2) \nonumber\\
                   &\propto \exp\left\{ -\frac{(s-m)^2}{\sigma^2}\right\} \exp\left\{ -\frac{m^2}{\sigma^2}\right\} \nonumber\\
                   &=\N\left(M\mid s/2,\sigma^2/4 \right)
\label{math:c1_mp} 
\end{alignat} 
\noindent Then using the following identity \citep{mixes}: 
\begin{equation} \int \mathcal{N}( x \mid \alpha z, \sigma_1^2 ) \mathcal{N}( z \mid \mu, \sigma_2^2 ) \ dz = \mathcal{N}(  x \mid \alpha\mu, \sigma_1^2+\alpha^2\sigma_2^2 ) \label{math:c1_id} \end{equation}  
We calculate the conditional sibling distribution similarly: 
\begin{alignat}{3}
&p(S_2 \mid S_1={s_1}) &&= \int p( S_2, M=m \mid S_1={s_1} ) \ dm && \nonumber \\[1.0ex]
& && = \int p( S_2 \mid M=m, S_1={s_1} ) p(M=m \mid S_1={s_1} ) \ dm && \nonumber \\[1.0ex]
& &&= \int p( S_2 \mid M=m ) p(M=m \mid S_1={s_1}) \ dm && \nonumber \\[1.0ex]
&  &&= \int\mathcal{N}\left(S_2\mid m, \frac{\sigma^2}{2}\right) \mathcal{N}\left(M=m\mid \frac{{s_1}}{2}, \frac{\sigma^2}{4} \right) dm && \nonumber \\[1.0ex]
& p(S_2 \mid S_1 = {s_1})  &&=\mathcal{N}\left(S_2\mid \frac{{s_1}}{2}, \frac{3\sigma^2}{4} \right) && 
\label{math:c1_sib} 
\end{alignat} 
\noindent Thus, as predicted by the \emph{infinitesimal polygenic model} the conditional sibling liability is normally distributed
around the midparent liability distribution with additional variance equal to half the population liability variance.
\newpage

%%%%%%%%%%%%%%%%%%%%%%%%%%%%%%%%%%%%%%%%%%%%%%%%%%%%%%%%%%%%%%%%%%%%%%%%%%%%% 
%%%%%%%%%%%%%%%%%%%%%%%  APPENDIX 1 - CASE 2  %%%%%%%%%%%%%%%%%%%%%%%%%%%%%%%
%%%%%%%%%%%%%%%%%%%%%%%%%%%%%%%%%%%%%%%%%%%%%%%%%%%%%%%%%%%%%%%%%%%%%%%%%%%%%
\subsection*{\normalsize Case 2) Index trait value known, continuous trait ($h^2\ne 1$): $P(S_2 \mid S_1 = s_1)$}   ~\\
In this case, the primary result considered in this manuscript, a
trait z-value, or equivalently, the percentile rank of an index
sibling in genome wide association where the rank-based inverse
transformation has been applied \citep{qtn} is known.  Transformation
to a $Z$ distribution ($\sigma^2=1$) means that for heritability $h^2$
the genetic liability and environmental contributions to trait
variance are are $\sigma^2_g=h^2$ and $\sigma_e^2=1-h^2$,
respectively. Similar to the previous case we begin by calculating the
conditional mid-parent liability from Bayes' theorem:
	
\begin{alignat}{2} 
p( M_g=m_g \mid S_1=s_1 ) &\propto p( S_1=s_1 \mid M_g=m_g ) p( M_g=m_g ) \nonumber \\
&= \N\left( S_1=s_1 \mid m_g, 1-\frac{h^2}{2} \right) \N( M_g=m_g \mid 0, h^2/2 ) \nonumber \\
&\propto\exp\left\{ -\frac{(s_1-m_g)^2}{2-h^2}\right\} \exp\left\{ - \frac{m_g^2}{h^2} \right\} \nonumber \\
&\propto\exp\left\{-\frac{2m_g^2-2m_gs_1h^2}{2h^2(1-h^2/2)}\right\}  \nonumber \\
&\propto\exp\left\{-\frac{(m_g-s_1h^2/2)^2}{h^2(1-h^2/2)}\right\} \nonumber \\
&=\N\left(M_g=m_g\mid\frac{1}{2}s_1h^2,\frac{1}{2}h^2(1-h^2/2) \right) 
\label{math:c2_mgGivenS} 
\end{alignat}

\noindent Then, we again use \eref{math:c1_id} to derive the distribution conditional sibling distribution:  
\begin{align}
p(S_2=s_s \mid S_1=s_1 ) &= \int p( S_2=s_2 \mid M_g=m_g ) p(M_g=m_g \mid S_1=s_1) \ dm_g \nonumber\\
&=\int\N\left( S_2\mid m_g,1-\frac{h^2}{2}\right) \N\left(M_g\mid\frac{1}{2}s_1h^2, \frac{1}{2}h^2\left(1-\frac{h^2}{2} \right)\right)  dm_g\nonumber\\
&=\N\left(S_2=s_2 \mid \frac{1}{2}s_1h^2,1-\frac{h^4}{4}\right) \label{math:c2_s2Givens1}
\end{align}
\newpage

%%%%%%%%%%%%%%%%%%%%%%%%%%%%%%%%%%%%%%%%%%%%%%%%%%%%%%%%%%%%%%%%%%%%%%%%%%%%% 
%%%%%%%%%%%%%%%%%%%%%%%  APPENDIX 1 - CASE 3  %%%%%%%%%%%%%%%%%%%%%%%%%%%%%%%
%%%%%%%%%%%%%%%%%%%%%%%%%%%%%%%%%%%%%%%%%%%%%%%%%%%%%%%%%%%%%%%%%%%%%%%%%%%%%
\subsection*{\normalsize Case 3) Multiple index trait values known, continuous trait: $P(S_3 \mid S_1 = s_1, S_2 = s_2)$} ~\\ 
The conditional sibling distribution can also be derived using the joint trait
distribution for related individuals:
\begin{align*}
y&\sim\N(0,\mathbf{G})\\
\intertext{where the covariance $\mathbf{G}$ is the genetic relationship matrix. Thus for a sibling pair}
p(s_1,s_2)&=\N_2\left(0, \left(\begin{array}{cc} 1 & h^2/2 \\
h^2/2 & 1
\end{array} \right)\right)
\end{align*}
As shown by Bernardo and Smith \citep{BandS}, if $X$ is multivariate
normal $\N(\mu,\lambda^{-1})$, where $\lambda=\Sigma^{-1}$ is the
precision matrix, and $X$ is partitioned into $x_1$ and $x_2$, with
corresponding partitions of $\mu$ and $\lambda$ of:
\[
\mu=\left(\begin{array}{c}\mu_1\\\mu_2\end{array}\right),\quad
\lambda=\left(\begin{array}{cc}\lambda_{11} & \lambda_{12}\\
\lambda_{21} & \lambda_{22} \end{array}\right)
\]
then the conditional distribution of $x_1$ given $x_2$ is also normal
with mean and precision matrix:
\begin{equation}  \mu_1-\lambda_{11}^{-1}\lambda_{12}(x_2-\mu_2),\quad\lambda_{11}\label{math:c3_gauss} \end{equation}
\noindent Thus, given the joint distribution for three siblings: 
\begin{align*}
p(s_1,s_2,s_3)&=\N_3\left(0,
\left(\begin{array}{ccc}
1 & h^2/2 & h^2/2\\
h^2/2 & 1 & h^2/2 \\
h^2/2 & h^2/2 & 1
\end{array} \right)\right)\\
\intertext{The precision matrix $\lambda=\Sigma^{-1}$ is}
\lambda&= \frac{1}{2+h^2-h^4}\left(\begin{array}{ccc}
2+h^2 & -h^2 & -h^2 \\
-h^2 & 2+h^2 & -h^2 \\
-h^2 & -h^2 & 2+h^2\\
\end{array} \right)\\
\intertext{And we can calculate the conditional distribution for for two sibling using \eref{math:c3_gauss}}
p(S_3 \mid S_1=s_1,S_2=s_2) &= \N\left(S_3\mid
\frac{h^2}{2+h^2}(s_1+s_2),
1-\frac{h^4}{2+h^2}\right)
\end{align*}
\newpage

%%%%%%%%%%%%%%%%%%%%%%%%%%%%%%%%%%%%%%%%%%%%%%%%%%%%%%%%%%%%%%%%%%%%%%%%%%%%% 
%%%%%%%%%%%%%%%%%%%%%%%  APPENDIX 1 - CASE 4  %%%%%%%%%%%%%%%%%%%%%%%%%%%%%%%
%%%%%%%%%%%%%%%%%%%%%%%%%%%%%%%%%%%%%%%%%%%%%%%%%%%%%%%%%%%%%%%%%%%%%%%%%%%%%

\subsection*{\normalsize Case 4) Binary trait: $P(S_2 \mid S_1 = A\hspace{-0.1em}f\hspace{-0.3em}f\hspace{-0.2em}ected)$} ~\\
Here we again assume an underlying distribution that is
$\mathcal{N}(0,1)$ and made up of genetic and environmental
components.  However, we only know the index sibling's \emph{status},
which, as described under the liability threshold model
\citep{falconer1967}, is equivalent to conditioning on the event where
than index sibling's trait value is above or below a z-value threshold
$T$:

\begin{equation}
  p(S_2 \mid S_1{=}\hspace{0.1em}\mbox{Affected}) = p(S_2 \mid S_1 >
  T) \hspace{0.8em} \mbox{and}\hspace{0.8em} p(S_2 \mid
  S_1{=}\hspace{0.1em}\mbox{\stkout{Affected}}) = p(S_2 \mid S_1 <
  T) \label{math:c4_liability}
\end{equation} 

\noindent where $T=\Phi^{-1}(1-K)$, $\Phi^{-1}$ is the inverse normal
cumulative distribution function, and $K$ is the incidence of the
binary trait in the population.  Thus, the conditional distribution
one sibling given an \emph{Affected} index sibling can be can be
calculated integrating over the normal distribution truncated at $T$:

\begin{equation} p(S_2 \mid S_1 > T) = \int_T^\infty p(S_2 \mid S_1 = s_1) p(S_1 = s_1) ds_1 \label{math:c4_trunc} \end{equation} 

\noindent The first two moments of the this truncated normal \citep{JKB} are: 
\begin{align*}
\E(S_1 \mid S_1>T ) &= \frac{\phi(T)}{1-\Phi(T)} = \frac{\phi(T)}{K} = i \\
\V(S_1 \mid S_1>T ) &= 1+T\frac{\phi(T)}{K}-\left(\frac{\phi(T)}{K}\right)^2 = 1+iT-i^2
\end{align*}
Approximating the index sibling distribution using a normal whose moments are taken from this truncated distribution, \eref{math:c4_trunc} becomes: 
\begin{align}
p(S_2=s_s \mid S_1 > T ) &= \int\N\left(S_2=s_2 \mid \frac{1}{2}s_1^*h^2, 1-\frac{h^4}{4}\right) \N(S_1=s_1^*\mid i, 1+iT-i^2 ) ds_1^*\\
\intertext{which can be solved using the identity given in \eref{math:c1_id}:}
&=\N\left(S_2=s_2\mid\frac{1}{2}ih^2, 1-\frac{h^4}{4}+\frac{1}{4}h^4(1+iT-i^2)\right) \nonumber \\
&=\N\left(S_2=s_2\mid\frac{1}{2}ih^2, 1-\frac{h^4}{4}i(i-T)\right) \label{math:c4_reich}
\end{align}
Thus, conditional on an \emph{Affected} sibling, the probability of concordance is: 
\begin{align*}
  p(S_2=\mbox{Affected}\mid S_1{=}\hspace{0.1em}\mbox{Affected}) &=p(S_2 > T \mid S_1 > T)\\
								 &=1 - \Phi\left(\frac{T-ih^2/2}{\sqrt{1-h^4i(i-T)/4}}\right)
\end{align*} 
which is equivalent to Reich's \citep{reich} correction to Falconer's
\citep{falconer1965} approximation where the relationship between the
relatives is 0.5 for siblings.  The probability of discordance given an \emph{Unaffected} sibling
can be calculated from Bayes' Theorem:
\begin{align*}
  p(S_2 = \mbox{Affected} \mid S_1{=}\hspace{0.1em}\mbox{\stkout{Affected}} ) &=
  \frac{p( S_1{=}\hspace{0.1em}\mbox{\stkout{Affected}} \mid S_2 = \mbox{Affected}) p(S_2=\mbox{Affected})}{p(S_1{=}\hspace{0.1em}\mbox{\stkout{Affected}})}\\
  &=\frac{K - K
    p(S_2{=}\mbox{Affected} \mid S_1{=}\mbox{Affected})}{1-K}
\end{align*} 
\noindent which allow the conditional probability of case status to be
determined given a index sibling's status.
\newpage
%%%%%%%%%%%%%%%%%%%%%%%  END APPENDIX-1 %%%%%%%%%%%%%%%%%%%%%%%%%%%%%%%%%%%%%%%%%%%%%%%%%%


%%%%%%%%%%%%%%%%%%%%%%%%%%%%%%%%%%%%%%%%%%%%%%%%%%%%%%%%%%%%%%%%%%%%%%%%%%%%%%%%%%%%%%%%%%%%%%%%%%%%%%%%%%%%%%%%%
%%%%%%%%%%%%%%%%%%%%%%%%%%%%%%%%%%%%%%%%%% APPENDIX2 %%%%%%%%%%%%%%%%%%%%%%%%%%%%%%%%%%%%%%%%%%%%%%%%%%%%%%%%%%%%
%%%%%%%%%%%%%%%%%%%%%%%%%%%%%%%%%%%%%%%%%%%%%%%%%%%%%%%%%%%%%%%%%%%%%%%%%%%%%%%%%%%%%%%%%%%%%%%%%%%%%%%%%%%%%%%%%
\section*{Appendix 2: Statistical Tests for Complex Architecture} 
\label{app:part2}
Here we describe our statistical tests for complex tail architecture.
Our tests identify changes in the conditional sibling distribution
when ascertaining on an index sibling in the tail relative to
polygenic expectation.  To carry out these tests we establish a null
distribution built on the assumption that indexing on siblings not in
the tails reduces that likelihood that either sibling phenotype in the
pair is driven by rare variants of large effect.  We use the region
from the $5^{th}$ to the $95^{th}$ percentile to estimate
heritability.  From the $n$ sibling pairs where the index sibling
$s^{(i)}_1$ is in the $5^{th}$ to the $95^{th}$, we calculate the
conditional likelihood
\eref{math:c2_s2Givens1}:
\begin{equation}
  \mathcal{L}(D | h^2)=\prod\limits_{i}^{n}p(s^{(i)}_{2} | s^{(i)}_{1},h^2) \propto
  \left(1-\frac{h^4}{4}\right)^{-n/2} \exp{ \left( -\frac{
        \sum\limits_{i}^{n}   \left(s^{(i)}_2 - \frac{s^{(i)}_1{h^2}}{2}\right)^2
      }{2\left(1-\frac{h^4}{4}\right)} \right) }
\end{equation} 
and maximize the log-likelihood with respect to $h^2$: 	
\begin{equation} \log \mathcal{L} \propto -n\log\left(1 - \frac{h^4}{4}\right) - \frac{1}{1-\frac{h^4}{4}} \sum_{i=1}^n \left( s_{2}^{(i)} - \frac{s_1^{(i)} h^2}{2} \right)^2 
\label{math:loglike_sup} \end{equation}
to obtain an maximum likelihood estimate for $h^2$ that is used to
define the null distribution for our statistical tests.
	
%%%%%%%%%%%%%%%%%%%%%%%%%%%%%%%%%%%%%%%%%%%%%%%%%%%%%%%%%%%%%%%%%%%%%%%%%%%%%
%%%%%%%%%%%%%%%%%%%%%%%  APPENDIX 2 - CASE 1  %%%%%%%%%%%%%%%%%%%%%%%%%%%%%%%
%%%%%%%%%%%%%%%%%%%%%%%%%%%%%%%%%%%%%%%%%%%%%%%%%%%%%%%%%%%%%%%%%%%%%%%%%%%%%
\subsection*{Statistical Test for De Novo Architecture} ~\\ 

\noindent We identify \emph{de novo} mutations of large effect by testing for
discordance between siblings relative to the polygenic null using the
conditional distribution of a sibling given index sibling from
(\eref{math:c2_s2Givens1}).  Since \emph{de novo} mutations typically result
in trait values in the tail of the distribution the test conditions on
those index siblings in a specified upper quantile $q$ of the
distribution, i.e. those sibling pairs $(s_1^{(i)},s_2^{(i)})$ such that
$s_1^{(i)}>\Phi(q)$, defined as the set $A_q$. We introduce an
additional parameter $\alpha$ where values of $\alpha < 0$ in the
right tail and $\alpha > 0$ in the left tail are indicative of
discordant siblings with trait values closer to the mean, giving a
log-likelihood:
\begin{align}
\log \mathcal{L} &= -\frac{1}{2(1-\frac{h^4}{4})} \sum_{i\in A_q}\left(s_2^{(i)}-\left(\frac{1}{2}s_1^{(i)}h^2+\alpha)\right)\right)^2\\
\intertext{The null hypothesis $H_0:\alpha=0$ is tested via a score test:}
U=\frac{d}{d\alpha}\log \mathcal{L} &= \frac{1}{1-\frac{h^4}{4}} \sum_{i\in A}s_2^{(i)}-\left(\frac{1}{2}s_1^{(i)}h^2+\alpha\right)\label{U-denovo} \\
I=\frac{d^2}{d\alpha^2}\log \mathcal{L} &= -\frac{n}{1-\frac{h^4}{4}}\label{I-denovo}\\
\intertext{And the score test for $H_0:\alpha=0$:}
z=\frac{U}{\sqrt{-I}}&=\frac{\sum_{i\in A_q}s_2^{(i)}-\frac{1}{2}s_1^{(i)}h^2} {\sqrt{n(1-\frac{h^4}{4})}}
\end{align}
\newpage
%%%%%%%%%%%%%%%%%%%%%%%%%%%%%%%%%%%%%%%%%%%%%%%%%%%%%%%%%%%%%%%%%%%%%%%%%%%%%
%%%%%%%%%%%%%%%%%%%%%%%  APPENDIX 2 - CASE 2  %%%%%%%%%%%%%%%%%%%%%%%%%%%%%%%
%%%%%%%%%%%%%%%%%%%%%%%%%%%%%%%%%%%%%%%%%%%%%%%%%%%%%%%%%%%%%%%%%%%%%%%%%%%%%
\subsection*{Statistical Test for Mendelian Architecture} ~\\ 

\noindent Here we test for excess concordance between siblings in the tails of the
distribution by testing for an excess number of observed siblings in
the tail $S_2>\Phi^{-1}(q)$ given the index sibling is in the tail
$S_1>\Phi^{-1}(q)$, where $q$ is the quantile of interest. Denoting
the set of index siblings in the tail by $A_q$ and the size of the set by
$n$, under the null of pologenicity we calculate the probability that
the conditional sibling exceeds $\Phi^{-1}(q)$ from the normal cdf and
compute the mean to define mean concordance under polygenicity $\pi_0$: 
\begin{align*}
  \pi_0&=\frac{1}{n}\sum_{i\in A_q} P\left(S_2^{(i)}>\Phi^{-1}(q)\mid S_1^{(i)}=s_1^{(i)}\right)\\
       &=\frac{1}{n}\sum_{i\in A_q} \left( 1 - \Phi \left(
         \frac{\Phi^{-1}(q)  - \frac{s_1^{(i)}
         h^2}{2}}{\sqrt{1-\frac{h^4}{4}}} \right)\right)
\end{align*}
Denoting the observed concordance (number of sibling pairs both
$>\Phi^{-1}(q)$) by $r$, the binomial log-likelihood (ignoring the
constant) is: 
\begin{align*}
\log \mathcal{L} &= r\log\pi + (n-r)\log(1-\pi)\\
U=\frac{d}{d\pi}\log L &= \frac{r}{\pi} - \frac{n-r}{1-\pi}\\
&=\frac{r(1-\pi)-(n-r)\pi}{\pi(1-\pi)}\\
&=\frac{r-n\pi}{\pi(1-\pi)}\\
I=\frac{d^2}{d\pi^2}\log L &= -\frac{r}{\pi^2} - \frac{n-r}{(1-\pi)^2}\\
&=-\frac{r(1-\pi)^2+(n-r)\pi^2}{\pi^2(1-\pi)^2}\\
\intertext{Assuming $r=n\pi$ such that $I$ is not a function of any particular observation:}
&=-\frac{n}{\pi(1-\pi)}
\intertext{And the score test for $H_0:\pi=\pi_0$:} 
z=\frac{U}{\sqrt{-I}}&=\frac{r-n\pi_0}{\sqrt{n\pi_0(1-\pi_0)}}
\end{align*}

\subsection*{Statistical Test for Non-Polygenic Architecture} ~\\                                                                                                                                                           
                                                                                                                                                                                                                            
\noindent General departures from polygenicity can be identified based on
the degree of discordance in conditional sibling distribution from the expectation. 
Assuming polygenicity, given index and conditional
sibling pairs $(s_1^{(i)},s_2^{(i)})$, from \eref{math:S2givenS1} we can
write
\begin{align*}
z_2^{(i)} = \frac{(s_2^{(i)}-\frac{1}{2}s_1^{(i)}h^2)}{
\sqrt{1-\frac{h^4}{4}}}\sim \N(0,1) \\
\end{align*}                      
\noindent Therefore, departures in polygenicity can be tested in trait
quantiles by testing the observed distribution of $z_2^{(i)}, i\in A_q$, where
$A_q$ is the set of index siblings $s_1^{(i)}$ in the quantile of
interest, relative to a standard normal distribution via the Kolmogorov-Smirnov test.
\newpage
%%%%%%%%%%%%%%%%%%%%%%%  END CASE 2 %%%%%%%%%%%%%%%%%%%%%%%%%%%%%%%%%%%%%%%%%%%%%%%%%%
%%%%%%%%%%%%%%%%%%%%%%%  END APPENDIX-2 %%%%%%%%%%%%%%%%%%%%%%%%%%%%%%%%%%%%%%%%%%%%%%%%%%

% Jensen
%%%%%%%%%%%%%%%%%%%%%%%%%%%%%%%%%%%%%%%%%%%%%%%%%%%%%%%%%%%%%%%%%%%%%%%%%%%%%%%%%%%%%%%%
%%%%%%%%%%%%%%%%%%%%%%%%%%%%%%%%%%%%%%%%%% APPENDIX3 %%%%%%%%%%%%%%%%%%%%%%%%%%%%%%%%%%%
%%%%%%%%%%%%%%%%%%%%%%%%%%%%%%%%%%%%%%%%%%%%%%%%%%%%%%%%%%%%%%%%%%%%%%%%%%%%%%%%%%%%%%%%
\section*{Appendix 3: Model Evaluation} 
\label{app:part3}
%\vspace*{-5mm}
Here we compare our theoretical derivations (\eref{math:S2givenS1})
that rely on the \emph{infinitesimal polygenic model} \citep{barton2017} with simulated offspring data (\textbf{Figure 7}).  
We also compare our model to our empirical simulation (see \myemph{Model \& Methods}) that draws allele frequencies and effect
sizes from publicly available GWAS data \citep{neale} for two traits to produce parent and offspring genotype and 
genetic liability (equivalent to trait value when $h^2=1$) (\textbf{Figure 8}).


\begin{center}                                                                                                                              
\includegraphics[width=\linewidth,height=6cm]{../Figs/behavior}                                                                                 
\captionof{figure}{Theoretical and simulated conditional
expectation and variance in liability (z-score) and rank across
index sibling percentiles for conditional sibling, midparents
and index siblings. Simulation drew one-thousand parent
liability values from $\mathcal{N}(0,1)$, these
were randomly paired to produce to midparents with liability
$m_i$, two offspring were subsequently drawn from
($\mathcal{N}(m_i, \frac{1}{2})$) and randomly assigned as
index and conditional siblings.}
\end{center}                   
	
	
		
\begin{center}                                                                                                                              
\includegraphics[width=\linewidth,height=6cm]{../Figs/simscore}                                                                                 
\captionof{figure}{For both \emph{Fluid Intelligence} and \emph{Standing Height} GWAS variants (on chromosome one) 
were used to simulate parent and offspring genotypes and liability values. 
Plots show that for both traits the offspring distribution is normal and that the sibling distribution is multivariate normal, 
in line with our theoretical prediction}. 
\end{center} 
\vspace*{2mm} 
These tests demonstrate that our theoretical framework accurately
reflects an additive polygenic trait in an outcrossing population.
Additionally these results demonstrate that deviations in the
conditional sibling distribution can be interpreted as non-polygenic
architecture or quantile specific environmental effects.
\newpage

%%%%%%%%%%%%%%%%%%%%%%%  END APPENDIX-3 %%%%%%%%%%%%%%%%%%%%%%%%%%%%%%%%%%%%%%%%%%%%%%%%%%

% Jensen  outline
%%%%%%%%%%%%%%%%%%%%%%%%%%%%%%%%%%%%%%%%%%%%%%%%%%%%%%%%%%%%%%%%%%%%%%%%%%%%%%%%%%%%%%%%%%%%%%%%%%%%%%%%%%%%%%%%%
%%%%%%%%%%%%%%%%%%%%%%%%%%%%%%%%%%%%%%%%%% APPENDIX3 %%%%%%%%%%%%%%%%%%%%%%%%%%%%%%%%%%%%%%%%%%%%%%%%%%%%%%%%%%%%
%%%%%%%%%%%%%%%%%%%%%%%%%%%%%%%%%%%%%%%%%%%%%%%%%%%%%%%%%%%%%%%%%%%%%%%%%%%%%%%%%%%%%%%%%%%%%%%%%%%%%%%%%%%%%%%%%
\section*{Appendix 4: Software Availability and Extended Results} 
\label{app:part4}
%\vspace*{-5mm}
We have made our code, sample data, and a brief tutorial available online at www.sibArc.net \citep{sibarc}.  
Below we display the summary statistics and the tail results for all eighteen UK Biobank traits analyzed and referenced in the manuscript.
\FloatBarrier 
\begin{figure}[h] \vspace*{-0.15cm}
%\begin{maybe}{showFig} \includegraphics[trim=0cm 0cm 0cm 0cm, clip, width=\linewidth]{../Figs/ukb3.png} \end{maybe} 
\begin{maybe}{showFig} \includegraphics[trim=0cm 0cm 0cm 0cm, clip, width=18cm]{../Figs/ukb3.png} \end{maybe} 
    \caption{\textbf{Application To UKB (Extended).}}
\label{fig:ukb} \end{figure} \FloatBarrier
%%%%%%%%%%%%%%%%%%%%%%%%%%%%%%%%%%%%%%%   UKB FIGURE  %%%%%%%%%%%%%%%%%%%%%%%%%%%%%%%%%%%%%%%%%%%%%%%%%%%%%%%%%%%%	

\end{document}
