## Supplementary figures and images for "Sibling Similarity Can Reveal Key Insights into Genetic Architecture"

### behavior.png

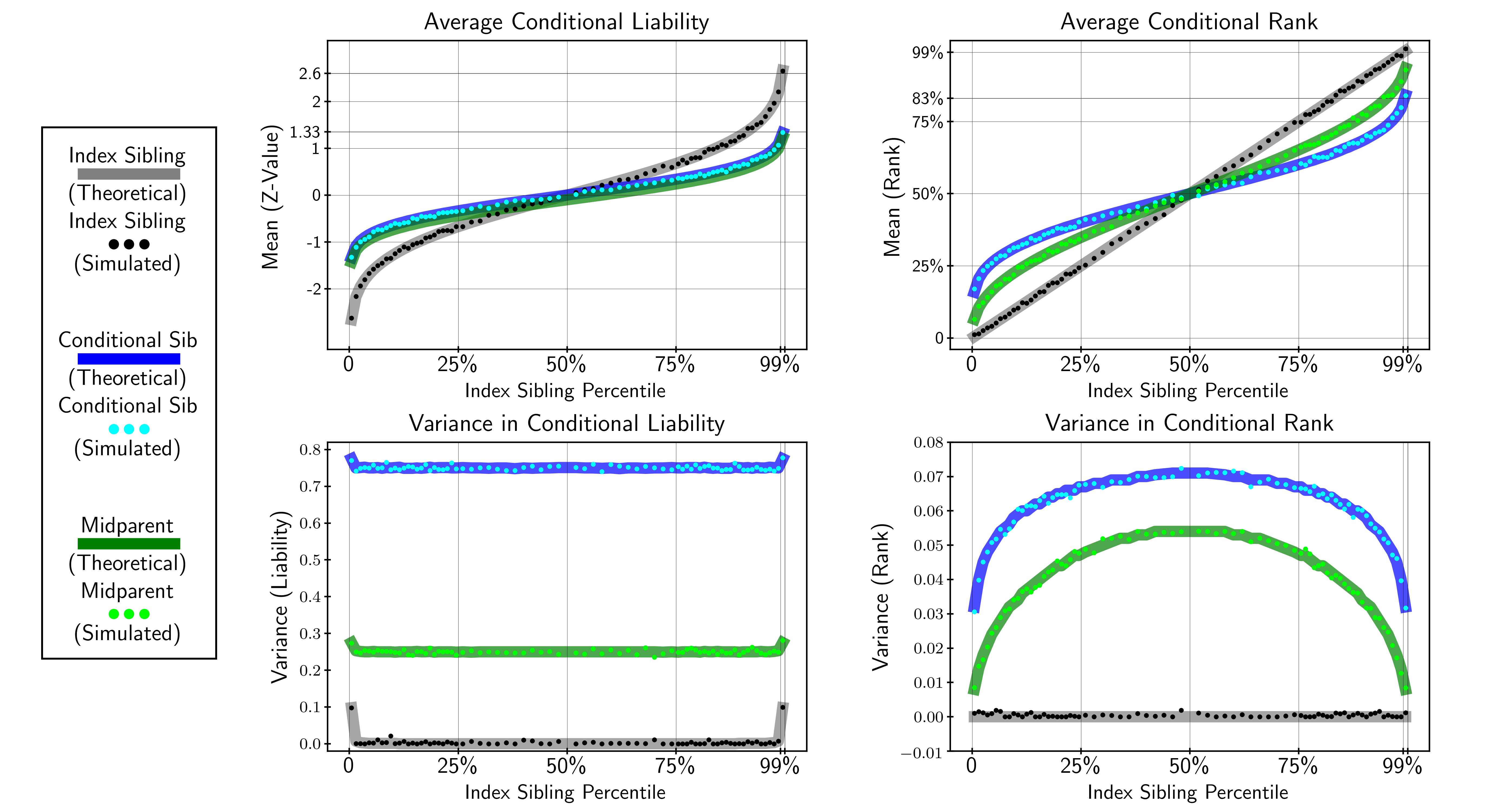

### intro.png

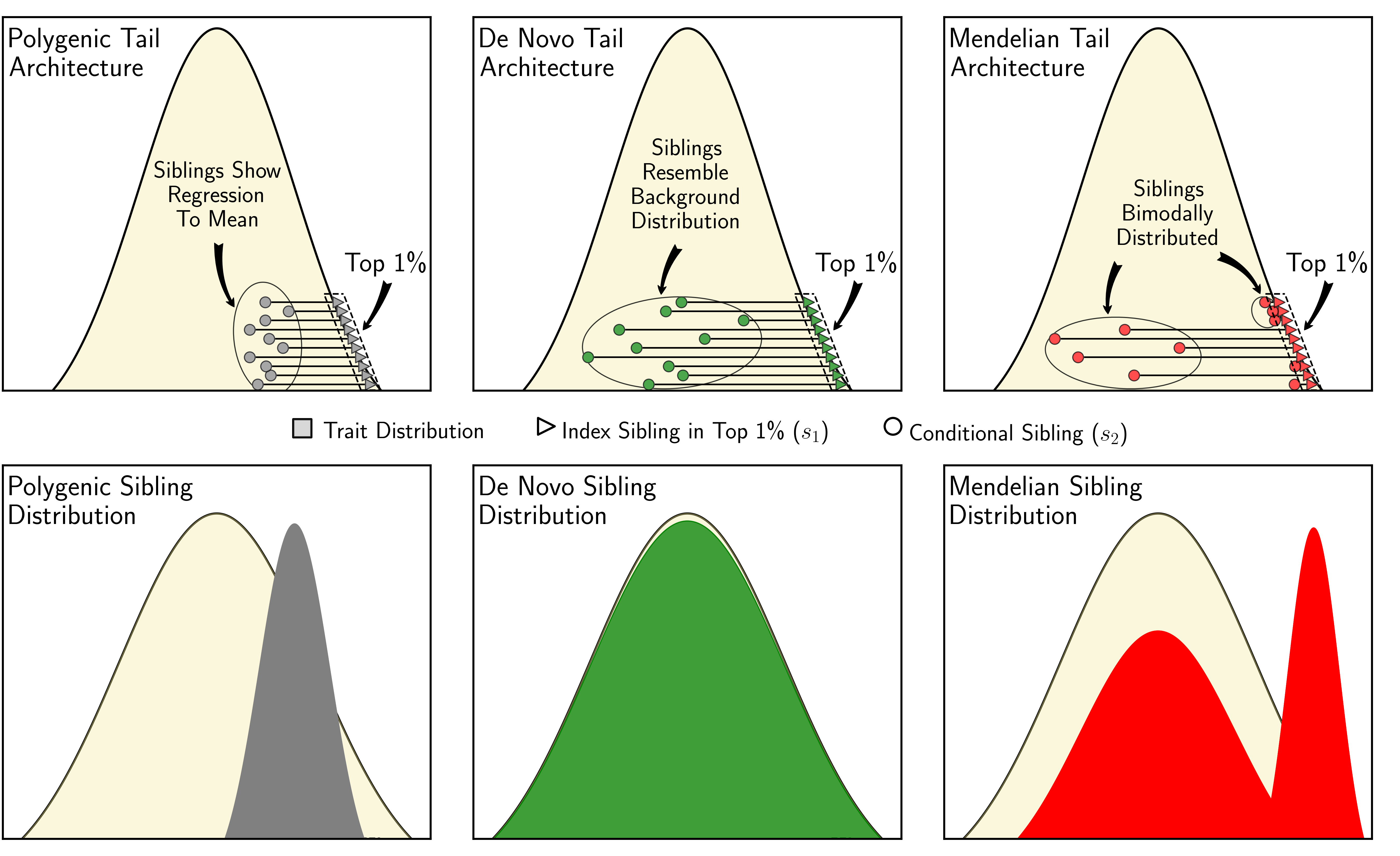

### schematic.png

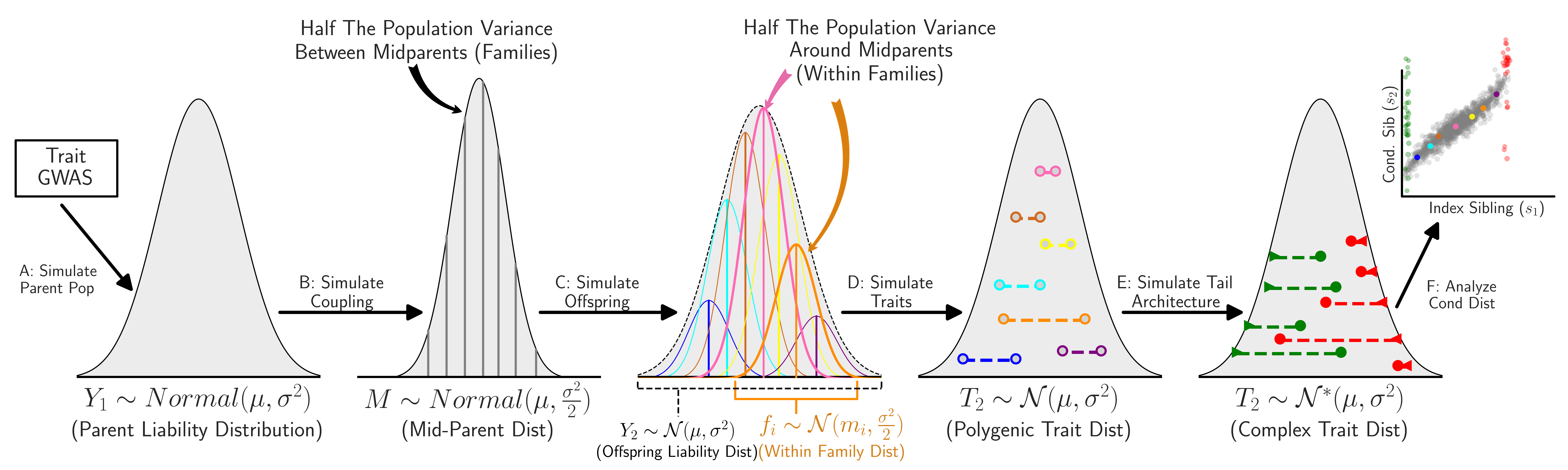

### sibdist.png

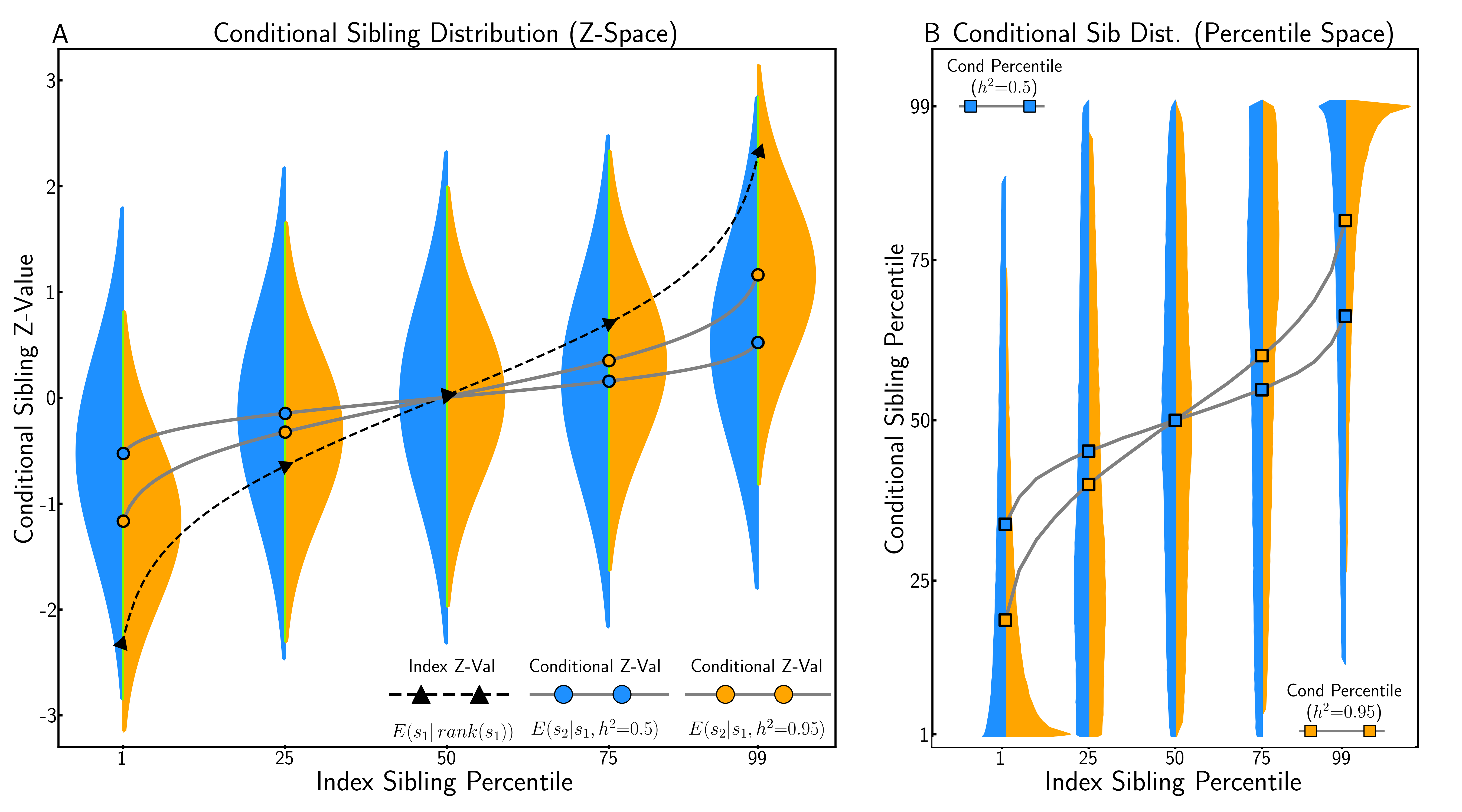

### side.png

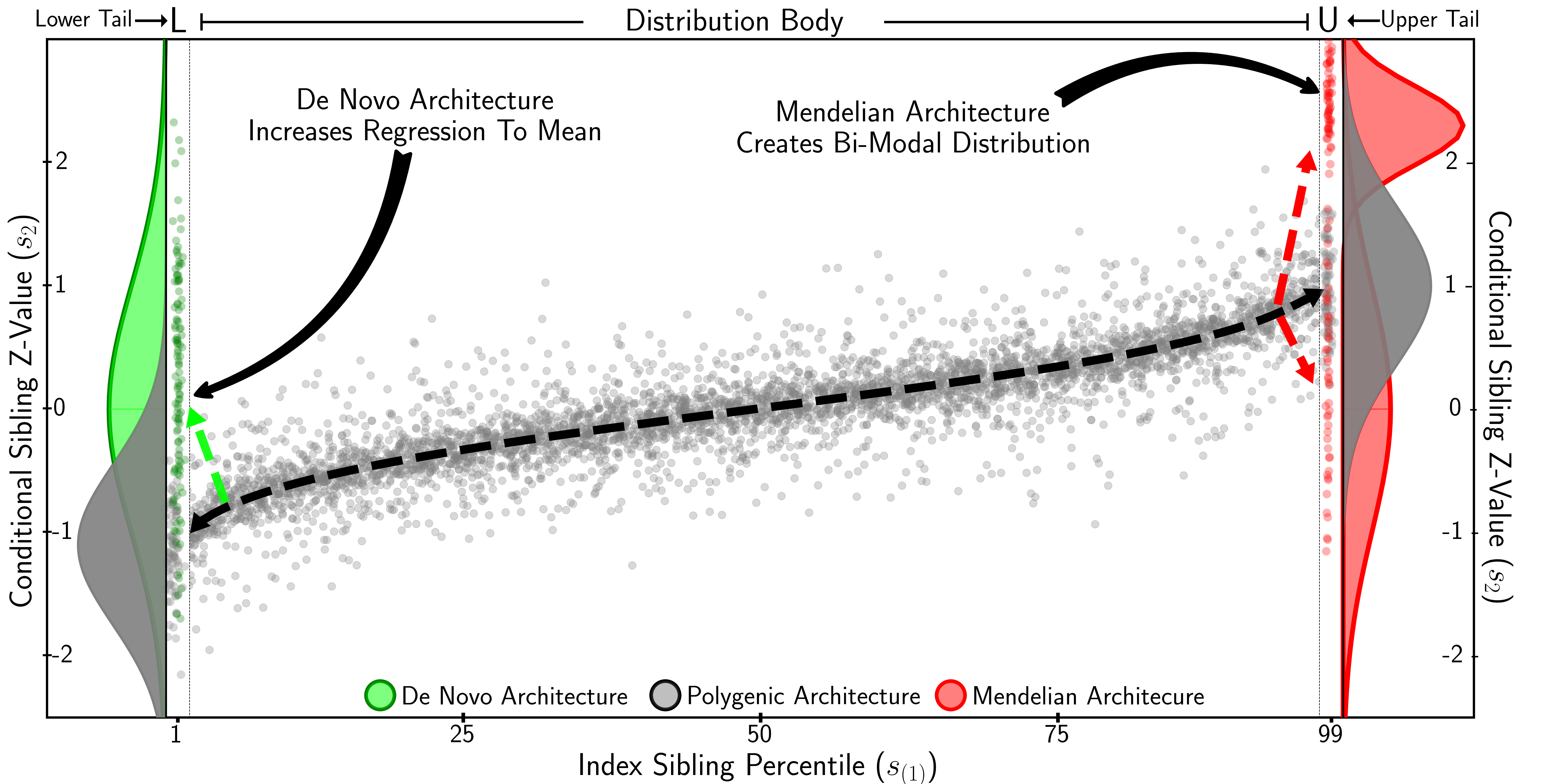

### simpower.png

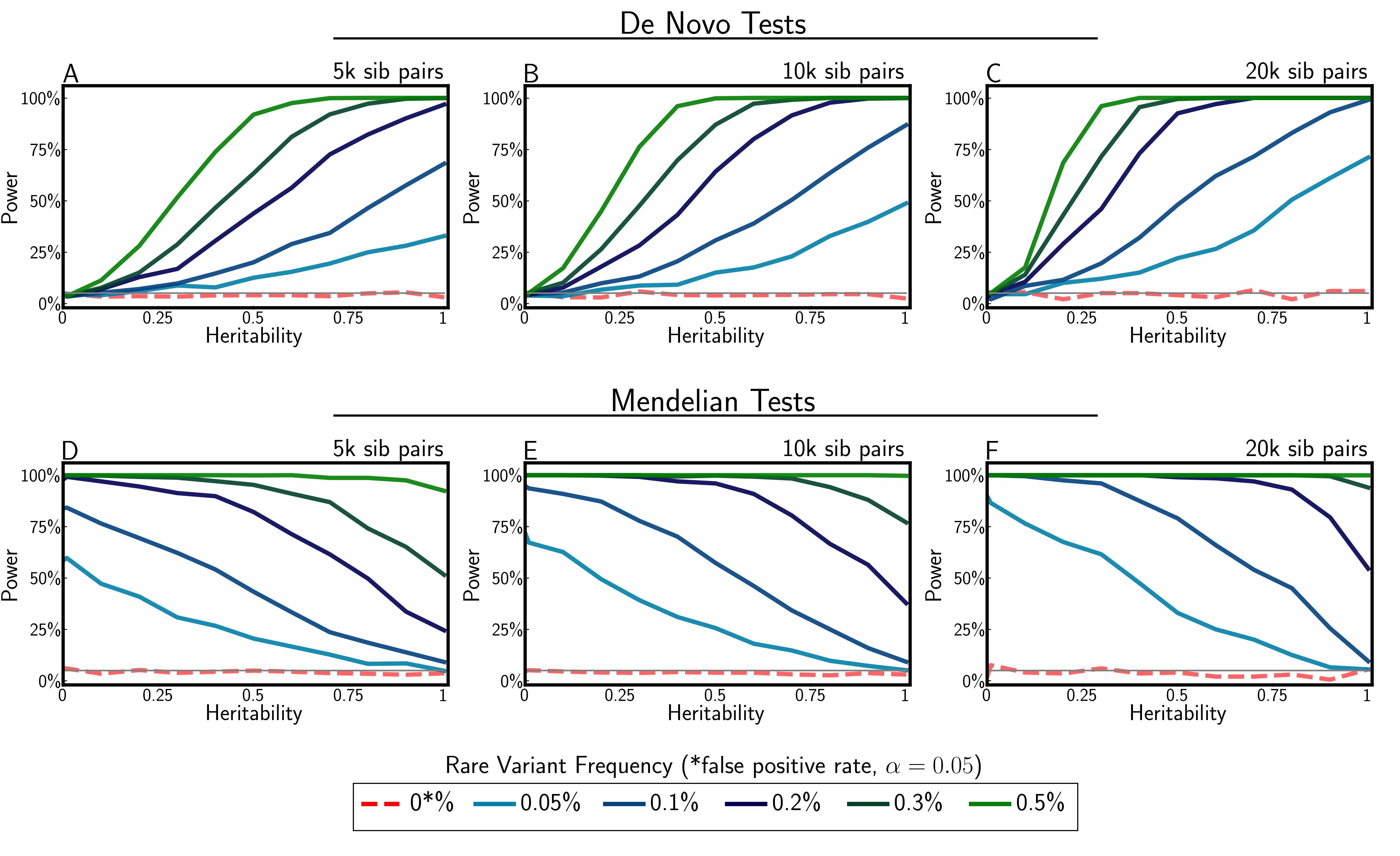

### simscore.png

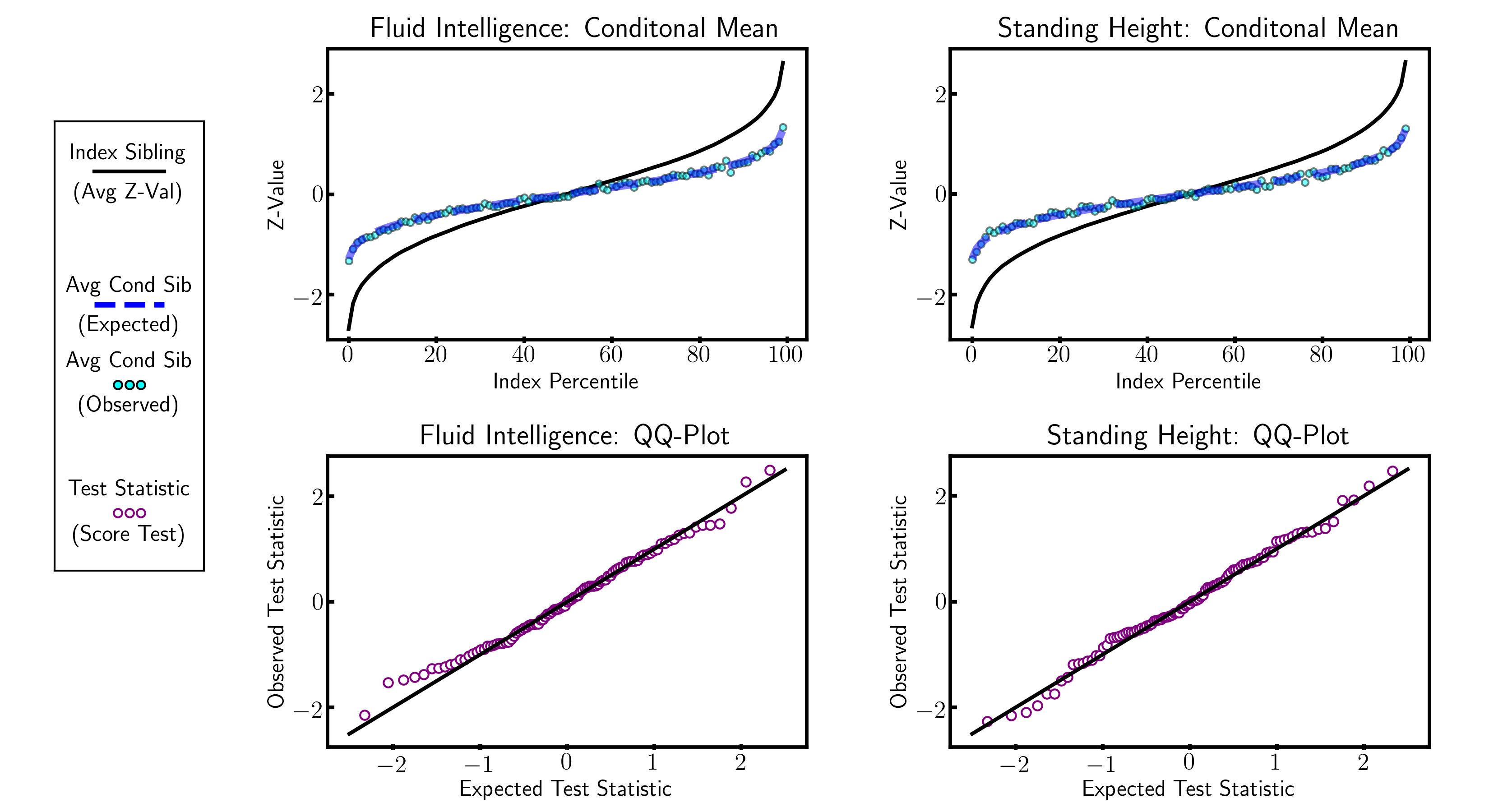

### ukb2.png

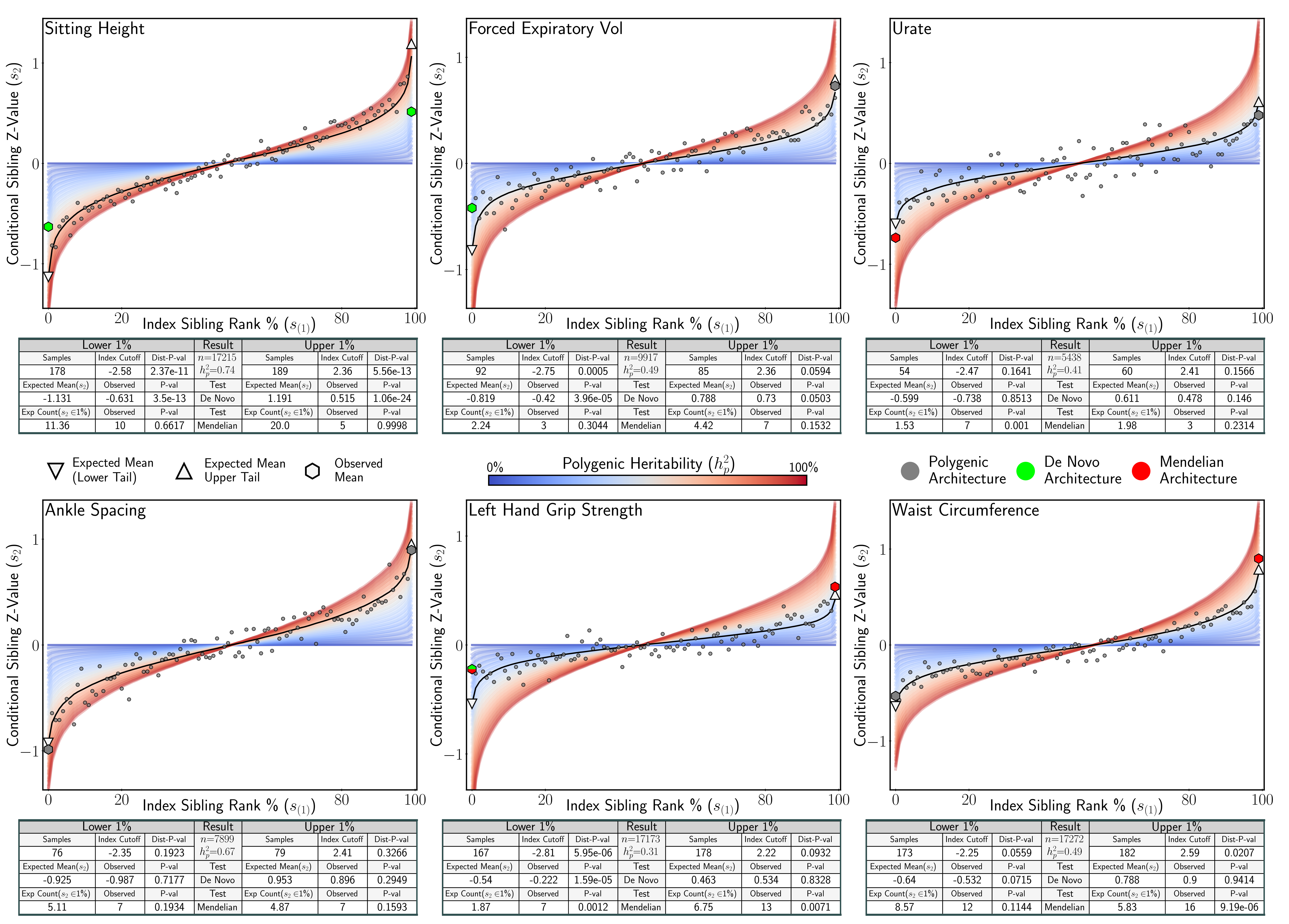

### ukb3.png

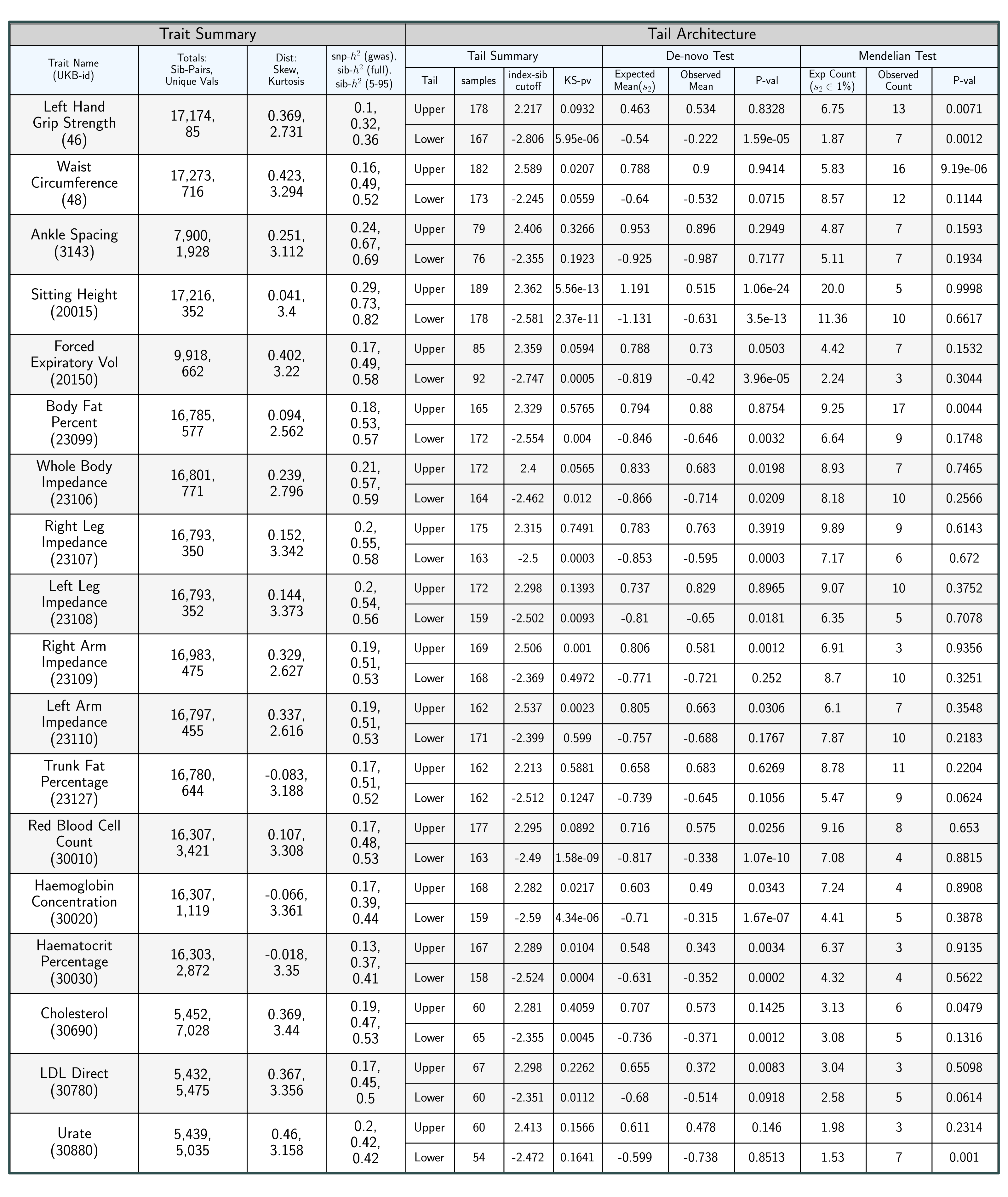
